# Sensor Movement Drives Emergent Attention and Scalability in Active Neural Cellular Automata

**DOI:** 10.1101/2024.12.06.627209

**Authors:** Mia-Katrin Kvalsund, Kai Olav Ellefsen, Kyrre Glette, Sidney Pontes-Filho, Mikkel Elle Lepperød

## Abstract

The brain’s distributed architecture has inspired numerous artificial intelligence (AI) systems, particularly through its neocortical organization. However, current AI approaches largely overlook a crucial aspect of biological intelligence: active sensing - the deliberate movement of sensory organs to explore the environment. To explore how sensor movement impacts behavior in image classification tasks, we introduce the Active Neural Cellular Automata (ANCA), a neocortex-inspired model with movable sensors. Active sensing naturally emerges in the ANCA, with belief-informed exploration and attentive behavior to salient information, without adding explicit attention mechanisms. Active sensing both simplifies classification tasks and leads to a highly scalable system. This enables ANCAs to be smaller than the image size without losing information and enables fault tolerance to damaged sensors. Overall, our work provides insight to how distributed architectures can interact with movement, opening new avenues for adaptive AI systems in embodied agents.

---

The neocortex plays a central role in perception, decision-making, language, and memory, exhibiting remarkable plasticity and resilience to damage. This has naturally inspired numerous AI systems’ architectures (Fukushima, 1980; Kohonen, 1990; George and Hawkins, 2009). However, vanishingly few AI models include the effect of sensor movement, ignoring that active sensing is a hallmark of perception in animals (Yang et al., 2016; Petersen, 2019; Wachowiak, 2011; Gilchrist, 2011; Nelson and MacIver, 2006). Unlike AI systems that statically classify images, active sensing allows attention to salient parts of the sensor space. As the AI community shows an increased interest in embodied AI, with AI for robotics and the proposal of embodied Turing tests (Lillicrap et al.), active sensing for AI becomes increasingly relevant as a more embodied approach to sensor-processing.

Sensor movement plays a crucial role in how organisms perceive and understand the world. For example, the somatosensory cortex does more than just process sensor input; it also projects to motor centers (Petersen, 2019; Rathelot et al., 2017). In the rodent whisker barrel cortex, a column (or barrel) has motor pathways to help move the whiskers during exploratory whisking (Petersen, 2019). Similar findings in the monkey (Rathelot et al., 2017) suggests that the somatosensory cortex is more frequently involved in movement than previously thought (Petersen, 2019). In the same vein, movement information is processed together with sensory information in sensory cortices (Kandel et al., 2021, p. 464, 583). Moreover, most senses are active. Common examples are olfaction (Wachowiak, 2011), and visual saccades, where the eyes flit between points of interest (Gilchrist, 2011). These two facts – that motor control is integrated in somatosensory cortex and that most sensing is active – has led researchers to reclaim the importance of sensor movement in perception and learning (Ahissar and Assa, 2016; Rao, 2024). Given these insights, we hypothesize that integrating movement in an artificial system might enhance its capabilities when engaging in sensor-processing.

Distributed architectures are common among systems inspired by the neocortex. One well-known example is Convolutional Neural Networks (CNNs), whose precursor (Fukushima, 1980, 1969) was inspired by the ocular dominance columns found in the visual cortex (Hubel and Wiesel, 1962). Similarly, Self-Organizing Maps (SOMs) (Kohonen, 1990), used for dimensionality reduction and clustering, drew inspiration from the somatosensory cortex in mammals. A more recent example, the Hierarchical Temporal Memory (HTM) model (George and Hawkins, 2009), uses a model of the canonical cortical column for various classification tasks and time series. These homogeneous distributed systems have several advantages. Firstly, parameter sharing allows for modular systems with fewer parameters to optimize, leading to more resource-friendly training. Homogeneous modular systems also typically have high redundancy and scalability, which improves robustness to partial damage.

Especially distributed architectures like the Neural Cellular Automata (NCA), or message passing ensembles in general, benefit from redundancy and lack of centralization (Mordvintsev et al., 2020; Mousavi et al., 2019; Pathak et al., 2019). NCAs use a single neural kernel to process every neighborhood of an image or grid over time, similar to a convolutional layer of a CNN. As opposed to CNNs, the output of the kernel at timestep *t* is never compressed with pooling or convolutions, but simply added to a computational substrate for the network to receive again at timestep *t* + 1 (Mordvintsev et al., 2020; Randazzo et al., 2020; Medvet et al., 2020; Pontes-Filho et al., 2022). Lacking centralization means that the architecture does not have a fixed size. Therefore, NCAs can change the size of the model after training, which is challenging with CNNs. Given its distributed architecture, with local information exchange and limited fields of view, NCAs are well suited to create AI models inspired by cortical columns.

In this work, we take inspiration from the whisker barrel cortex in rodents, commonly agreed to have both sensor movement and distributedness. The architecture of the barrel cortex is well studied: A sheet of neural modules called barrels or columns maps one-to-one with whiskers on the rodent’s snout (Woolsey and Van der Loos, 1970; Petersen, 2019; Kandel et al., 2021, p. 457). The barrels process the input from the whiskers and help control them and are sparsely connected to their neighboring barrels (Petersen, 2019). Given this architecture, the barrel cortex may be abstracted into a message-passing ensemble of independent modules. The simplicity of the proposed system gives us the ideal foundation for exploring the behavior of sensor movement in a distributed system. Given the possibility that movement becomes underdeveloped, we seek to investigate how and if the sensors move when trained on real data. Additionally, we seek to investigate if a moving system has any advantages or disadvantages compared to a non-moving system.

To answer these questions, we present the Active Neural Cellular Automata (ANCA): A novel architecture that expands on NCAs with dynamic control over their sensor fields, enabling active sensing behaviors as observed in the whisker barrel cortex. We show that the ANCA demonstrate emergent active attention, as sensor fields purposefully move and converge on salient points. This mechanism simplifies image classification by ignoring irrelevant parts of the task space. Moreover, its distributedness allows the ANCA to scale, and movement is shown to be crucial for maintaining the performance as the system scales. Additionally, the ANCA exhibits promising fault tolerance to system perturbation, which suggests the ANCA could be useful for embodied robots with distributed sensors. Taken together, our results suggest that attentive behavior can emerge in distributed sensorimotor systems as a direct consequence of their ability to move. Unlike NCAs with explicit attention modules (e.g., (Tesfaldet et al., 2022)), ANCAs attentive behavior arises without explicit attention modules. While being useful for object recognition and decision making, this key insight may also contribute to understanding how perception and attention could be linked in moving systems. This paves the way for further research into the benefits of distributed AI systems in robots equipped with movement.

## 1 Results

### 1.1 The Active Neural Cellular Automaton

In this work, we introduce the **Active NCA**, a novel system that builds on the established NCA to employ active sensing. This active behavior is achieved by allowing convolutional kernels to move independently within an input image, mimicking how whiskers can move over objects (see Figure 1 for an overview).

**Figure 1:**
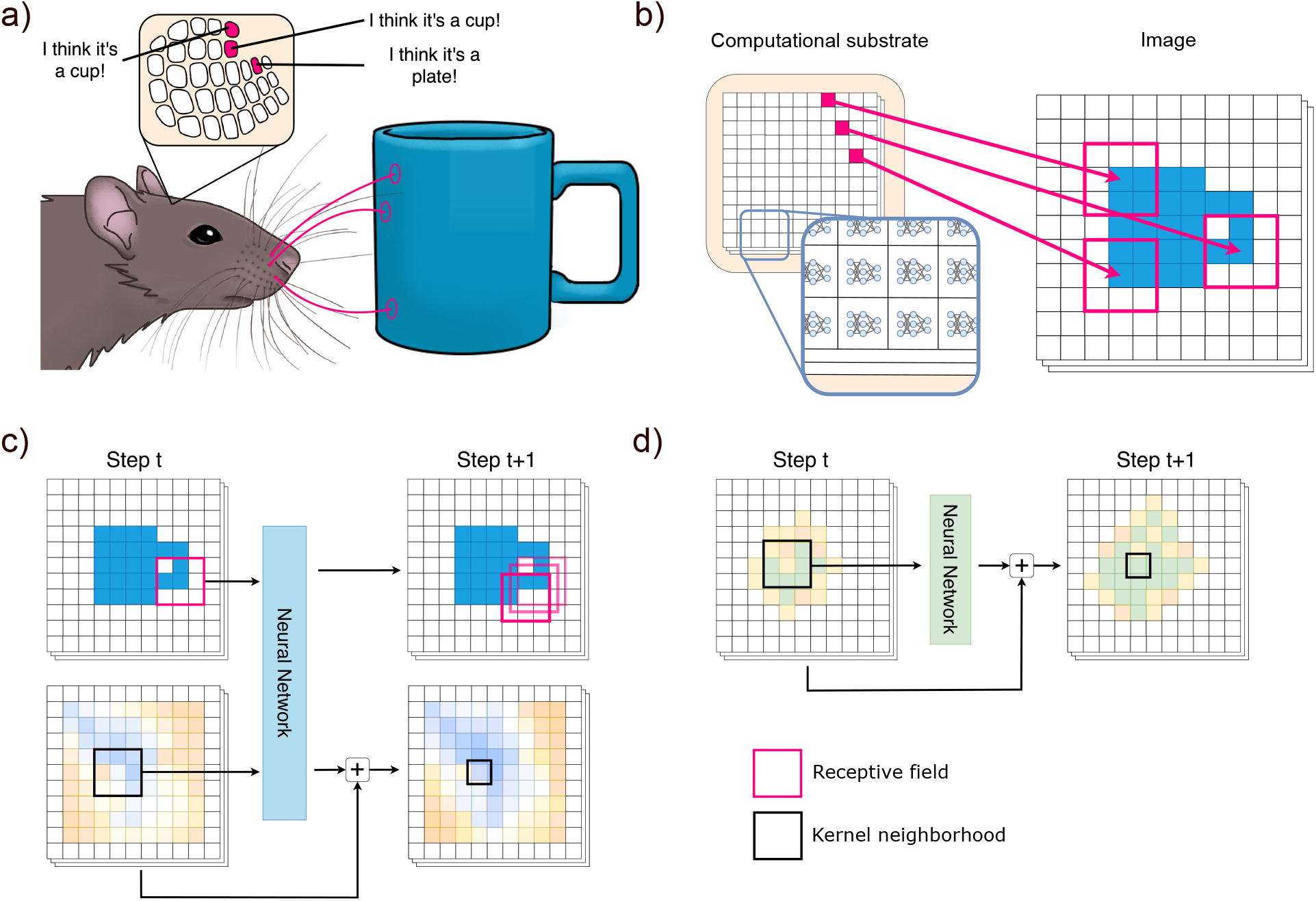
The Active Neural Cellular Automaton. a) The whisker barrel cortex (top) has columns/barrels corresponding to each whisker on the whisker pad. As whiskers move across objects, the barrels process the sensor input and form beliefs. b) We abstract the barrel cortex into an NCA substrate (left) that controls the receptive fields in the input space (right). c) The ANCA: For each kernel, its receptive field and substrate neighborhood is processed. The substrate then gets updated, and the receptive field is moved. d) The standard NCA: For each kernel, its substrate neighborhood is processed, and the substrate gets updated.

An NCA uses stationary kernels to process a substrate and update it over time (Randazzo et al., 2020; Mordvintsev et al., 2020). The substrate is a matrix of arbitrary size *N*_*neo*_ × *M*_*neo*_ × (*H* + *C*), where *N*_*neo*_ × *M*_*neo*_ is the size of the kernel grid, *H* represents the number of hidden channels for internal memory, and *C* channels for output. If classifying an image with an NCA (Randazzo et al., 2020; Walker et al., 2022), the substrate is the size of the image, and *C* corresponds to the number of class predictions. The substrate is similar to a hidden layer in a Recurrent Neural Network and can often be trained through backpropagation through time. Loss is computed pixel-wise across the *C* channels, first for every pixel, then averaged.

The ANCA diverges from NCAs in two ways:

1. We fully separate the image (the outside world) from the computational substrate (the cortex). The kernels are fixed in the substrate, but output actions to move their sensor (the whisker) in the image.
2. We supply each kernel with its current position in the image.

In the ANCA substrate, the kernels play the role of cortical columns, processing their sensor input (called the receptive field) and substrate neighborhood at time *t*, then moving their sensors and altering the substrate at time *t* + 1. The system operates for episodes of 50 timesteps. Receptive field size and substrate kernel size are independent, but here chosen to be 3 × 3 for both. As seen in section 1.4, kernels receiving their current position in the image improves performance. The code is found online 1.

The proposed Active NCA is non-differentiable, as the kernels must take actions to move their receptive fields. Therefore, the system was trained using the gradient-free optimizer Covariance Matrix Adaptation Evolutionary Strategy (CMA-ES) (Hansen and Ostermeier, 1996). We use Categorical Cross Entropy (CE) loss. For prediction, the class channels are averaged into one classification. Exact training details and metrics can be seen in section 3.2 and 3.3.

### 1.2 The system can classify complex data

The ANCA was tested on the standard machine learning benchmarks of MNIST (Deng, 2012), Fashion-MNIST (Xiao et al., 2017), and CIFAR-10 (Krizhevsky et al., 2009). With CIFAR-10, long training times from the CMA-ES optimization caused us to choose only 4 classes: Airplanes, ships, horses, and deer. The selected datasets provided increasing levels of complexity, enabling us to systematically evaluate the ANCA’s performance across different scenarios. With both simple objects, complex objects, and colored objects with specific contexts, this dataset selection helped demonstrate the ANCA’s versatility across varying levels of perceptual complexity.

While no extensive tuning was conducted, the networks were roughly hand-tuned. The resulting networks for MNIST and Fashion-MNIST were linear, with one hidden layer, while the CIFAR-10 network had 2 layers and Rectified Linear Unit (ReLU) activation (Zeiler et al., 2013). The output layer was always linear. A difference in system behavior between substrate sizes led us to collect networks with different substrate sizes as well. All scores are given in the appendix.

Establishing the system as a working classifier, we see that the ANCA performs moderately well across the datasets (Table 1), with the best accuracy on MNIST being 91.94%, Fashion-MNIST 76.04%, and CIFAR-10, 4 classes, 70.70%. In the literature, only Randazzo et al. (2020) have classified MNIST with an NCA, and they achieved a best accuracy of 96.2% during the episode and a best accuracy of 95.3% at the end of the episode. Comparatively, the ANCA could also have benefited from a larger network (6.6k parameters compared to the 22.5k of Randazzo et al.), but the training time with CMA-ES proved to be prohibitive. Overall, the method does learn to classify complex data, but more research would be needed to get more competitive scores.

**Table 1:** The performances of the ANCA on the datasets. The collected networks were tested on the train and test set. The best networks were found through lowest CE loss on the train set, and their best test-set scores are reported in the “best test” row. The mean scores average the performance of all the collected networks. These are the best results achieved with the ANCA. Further accuracies for different hyperparameters used in this paper can be seen in the appendix.

|  | MNIST 5 class | MNIST 10 class | Fashion-MNIST 10 class | CIFAR-10 4 class |
| --- | --- | --- | --- | --- |
| Best train | 96.34% | 91.57% | 76.45% | 69.37% |
| Best test | 97.27% | 91.97% | 76.04% | 70.70% |
| Mean train | 93.61±1.8% | 88.99±1.9% | 75.08±1.3% | 66.86±1.9 |
| Mean test | 94.50±1.8% | 89.53±1.7% | 74.36±1.4% | 68.13±1.8 |

### 1.3 Active sensing emerges from movement

Upon seeing how the ANCAs behaved, it became clear that viewing strategies would emerge. Therefore, we investigate their behavior considering a specific animal sensing strategy: Active sensing, a purposeful process that moves sensors to salient parts of the input space to form a belief through hypothesis-testing (Rao, 2024). Although no agreed-upon criteria of active sensing exist, we synthesize the current literature to identify three key aspects relevant to our system:

1. Focus: Information might be gained from “salient points”, as in foveation (Gilchrist, 2011; Yang et al., 2016).
2. The viewing path: Information might be gained from paths over time, as in whisking (Petersen, 2019; Nelson and MacIver, 2006; Hartmann, 2001).
3. Hypothesis-testing: Whether the ANCA’s belief (its current classification) inform its actions (Rao, 2024; Pezzulo et al., 2024).

To get an overview of the ANCA’s behavior, we plotted focus and the viewing path. For focus, the image neighborhoods occupied by receptive fields were recorded and averaged for all 50 timesteps. For the viewing path, kernel behaviors were averaged and displayed as colored arrows that showed their action and belief at every visited image coordinate. We also include videos where full episodes are displayed.

Figure 2 shows how the fields focus their receptive fields on salient parts of the input space, with edge-focusing behavior as a common denominator across datasets. On MNIST, ANCAs often target only the lower parts of the digits, not needing the whole picture for a correct classification. On Fashion-MNIST, focus varies: Sleeves, collars, or legs are targeted depending on the garment. For CIFAR-10, ANCAs focus on legs and bodies, but spend more time on the background. Likely, this reflects that when classes are similar (deer vs horses), the classes mainly differ in context.

**Figure 2:**
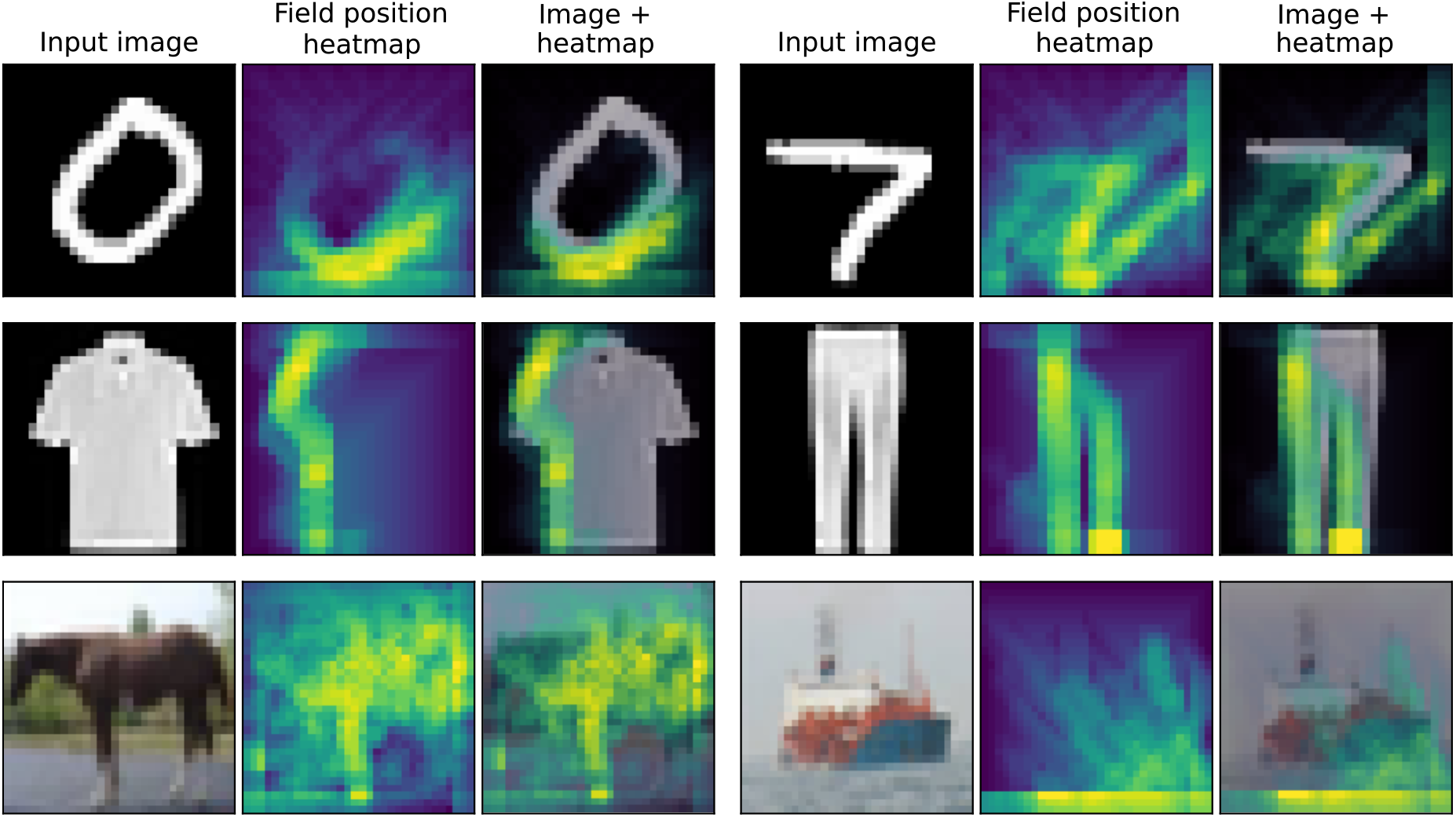
The model displays focusing behavior on salient points. Top: Focus of the best performer on MNIST (91.97% accuracy). Middle: Best performer of Fashion-MNIST on a 15 × 15 substrate (73.37% accuracy). Bottom: Best performer on CIFAR-10 4 classes (70.7% accuracy).

For temporal behavior, we show an ANCA classifying Fashion-MNIST images. It was trained on a larger substrate to make the behavior more visually clear. Figure 3a shows that the ANCA displays three distinct behaviors for this dataset: One for trousers, one for shoes, and one for torso garments. Within the groups shoes and torso garments, the behavior is very similar, as seen in figure 3b. The confusion matrix (figure E5, Appendix), reveals that misclassification is highest among classes that elicit the same behavior. Overall, the behavior of the system seems determined by what image is being viewed. We hypothesize that an initial sweep decides which group it is, prompting a distinct group sweep. This distinct sweep then refines the classification.

**Figure 3:**
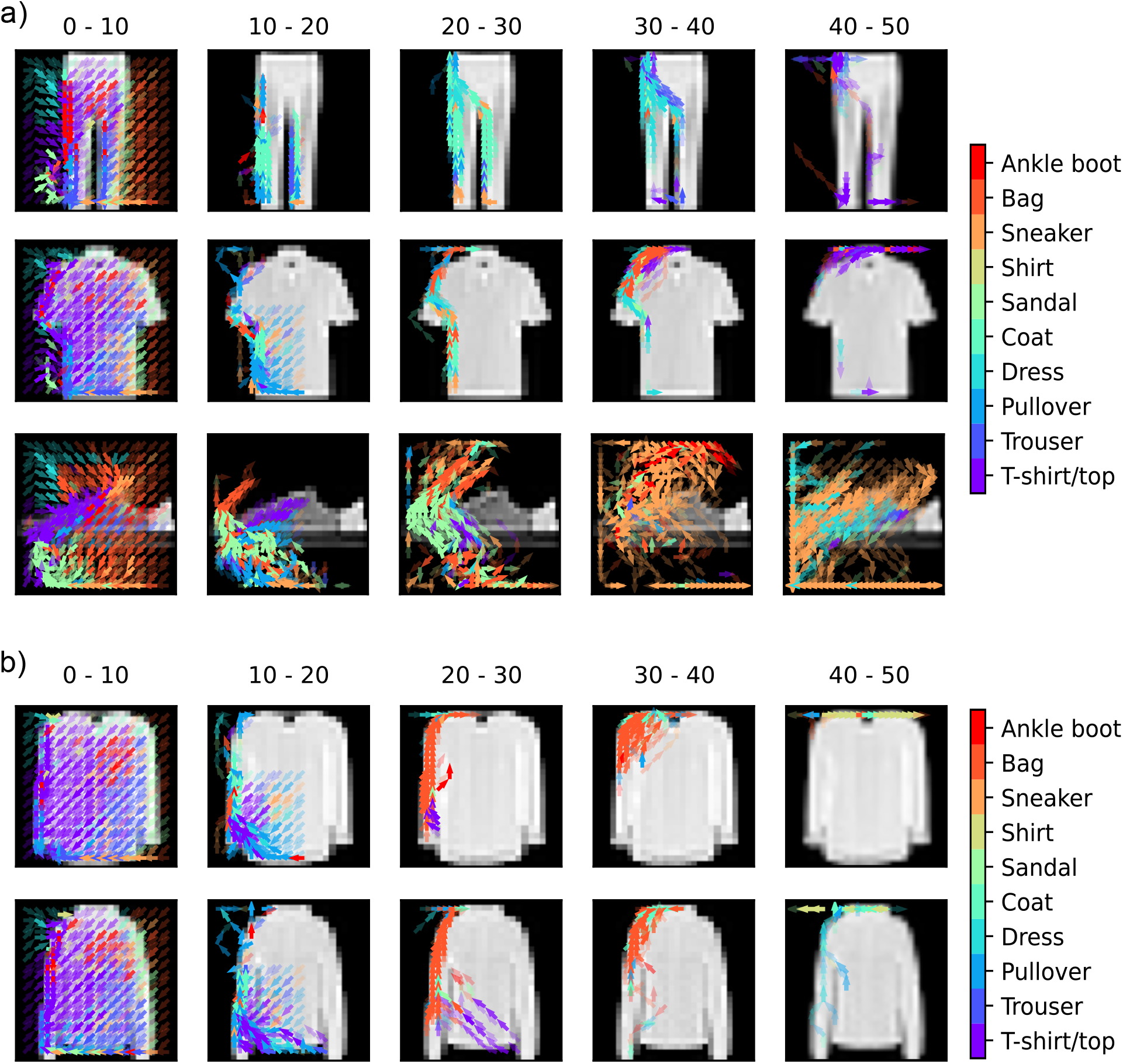
The viewing path on Fashion-MNIST. The ANCA displayed has a substrate of 15 × 15 and an accuracy of 73.37%. These images are all classified correctly. a) **Three distinct behaviors can be seen from the same ANCA**. Top: Trousers. Middle: T-shirt. Bottom: Sneaker. b) **Similar object elicit the same behavior**. Top: Shirt. Bottom: Coat.

When comparing small and large substrates (figure A1, Appendix), we find that larger substrates have more homogeneous behavior, while smaller substrates have more varied behavior. In short, smaller substrates rely more on sweeping behavior and following edges, while the larger substrates focus more on salient coordinates. Likely, this difference arises because more homogeneous behavior can emerge when only a low number of kernels receive 0-padded input. For small substrates, about 50% of kernels gets padded input compared to under 25% for large ones.

The ANCA’s classification behavior – moving over the image and focusing on salient points – demonstrates active sensing. This active sensing mechanism has a similar effect to a self-attention mechanism, where select parts of the input space is attended. However, the ANCA’s attentive behavior emerges completely without explicit attention modules or encouraging movement in the loss function. Next, we explore what benefits this attentive behavior can lead to in moving systems.

### 1.4 Movement can simplify the task and avoid irrelevant information

To gain insight into the effects of movement, we first applied it to a toy dataset: An image of a cup, knife, and bowl (Figure 4a). By increasing the space around the objects, we can increase the amount of irrelevant neighborhoods in the input, which complicates the classification task for distributed models. The dataset has no variation and no train and test set.

**Figure 4:**
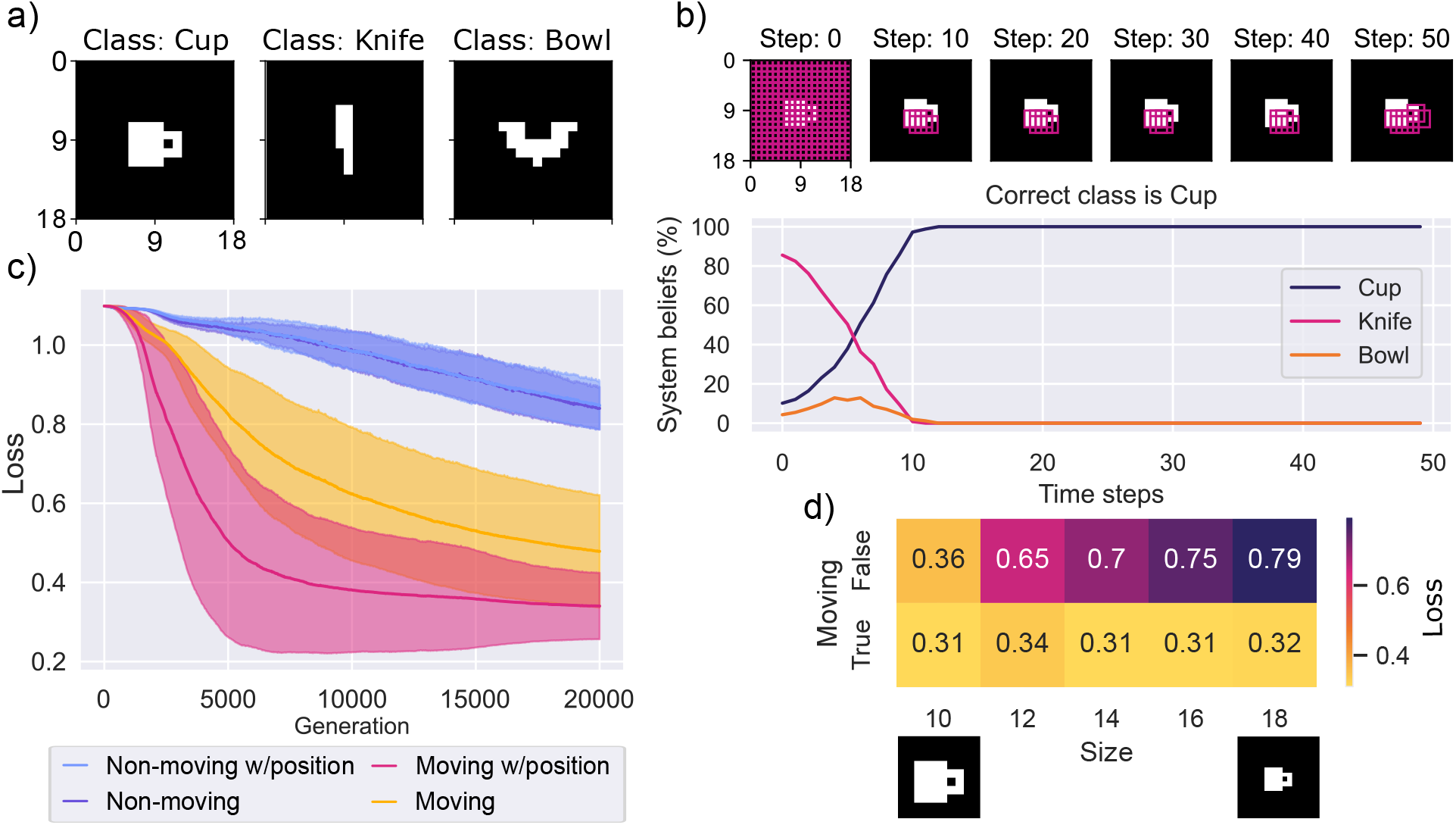
Movement helps avoid irrelevant information. a) A hand-made dataset of three objects with ample space around the objects. b) Behavior of the moving system with position information.c) The mean loss plots of each condition. The shaded areas are standard deviations. d) Testing the moving ANCA with position information and the non-moving ANCA without position information for increasing levels of empty space. The best hyperparameter pair was chosen from figure c, and 4 samples collected per size. For the loss values seen, the last 10 generations were averaged.

We performed an ablation study to explore the central features of the system: Movement and position information. This led to four system conditions to test: (1) Moving with position, (2) moving without position, (3) non-moving with position, and (4) non-moving without position. Further details in section 3.7.

Looking at figure 4c, the moving ANCAs learn the task faster than the non-moving ANCAs. We can easily see why: The non-moving ANCA cannot change its inputs and is forced to classify empty space. Testing the systems with increasing levels of space around the object confirmed this, as the non-moving system’s performance declines (Figure 4d). In contrast, the moving ANCA maintains performance by finding and focusing on the object (Figure 4b). This shows that the ANCA can ignore irrelevant regions of the image, as well as simplify tasks by focusing on important pixels. In addition, we see that position information can lead to faster convergence for the ANCA. With position information, the kernels find the object easier and can correlate pixel content with its position for an easier classification.

### 1.5 Movement makes the ANCA highly scalable

At some point in primate evolution, rapid expansion of the neocortical sheet increased cognitive capacity. One explanation for this growth, the radial unit hypothesis, suggests that neural progenitors give rise to cortical columns (Rakic, 2000), with each column adding potential cognitive capacity. As the primate brain has evolved to increase its neural progenitors (Kandel et al., 2021, p. 1141-1142), this columnar architecture appears highly scalable over evolutionary timescales.

Inspired by this, we tested the scalability of the ANCA. Unlike fixed architectures like CNNs, ANCAs’ distributed nature allows the computational substrate to be scaled independently of the image after training. However, the fact that it *can* scale up does not mean it will function well after scaling. Therefore, we trained the ANCA with position information – with and without movement – on six substrate sizes and tested their accuracy on sizes from 1 to 26× 26.

The ANCA (Figure 5b) has a clear advantage when it comes to scaling down compared to the nonmoving ANCA (Figure 5a). When not moving, the receptive fields cannot cover the full image for any substrate size smaller than the image. Because of this, information can be lost. Comparatively, the moving ANCA can always cover the entire image, given sufficient timesteps. Although in this simple experiment, the non-moving ANCA reached good accuracy for small substrate sizes, in general only the moving ANCA can guarantee good performance at any size. In practical terms, this allows the ANCA to be trained on smaller substrate sizes, reducing training time.

**Figure 5:**
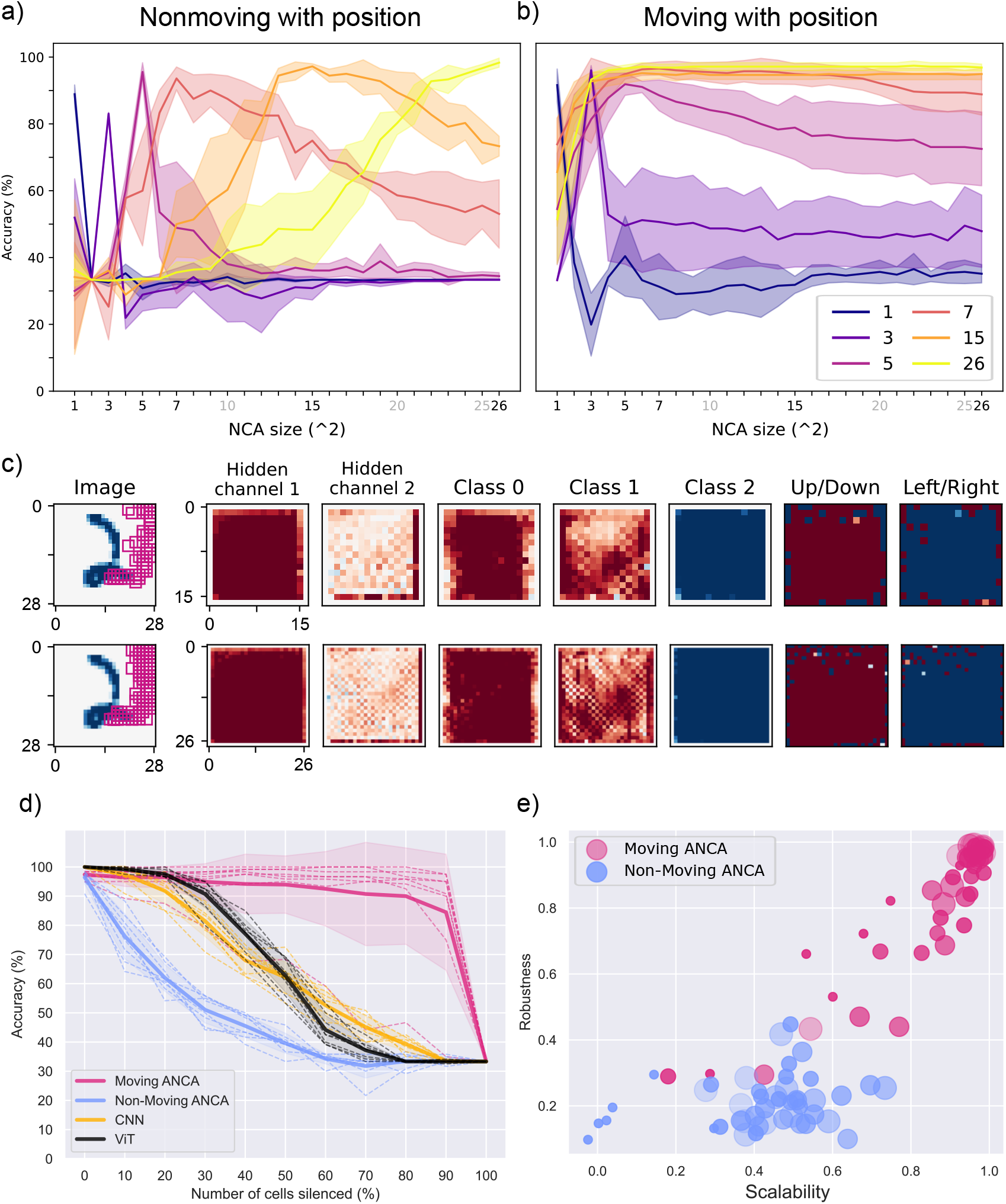
The scalability and fault tolerance of the ANCA. Pictured is the measured scalability of the non-moving (a) and the moving ANCA (b) for substrate sizes *x × x* where *x* is 1, 3, 5, 7, 15, and 26. Along the horizontal axis is the substrate test size, and the vertical axis displays the recorded accuracy at this test size. All systems were trained on 3 classes of MNIST, and as such, 33.33% is the lowest accuracy. c) Top: A trained ANCA on 0-2 digits from MNIST was recorded. The image is shown with field positions as pink squares. The substrate is shown with −1 to 1 as red to blue. The action outputs are shown with −0.0007 to 0.0007 as red to blue. Bottom: The same ANCA was tested on the same digit 2 with an expanded substrate (26 ×26). Overall values are maintained between the two substrate sizes: Notice how all colors are the same. The receptive field positions also maintain their behavior and end up in the same positions. d) The recorded accuracy (vertical axis) of an ANCA with position, non-moving ANCA with position, a CNN, and a ViT. Each model was recorded for increasing levels of silenced columns/patches (horizontal axis). e) The fault tolerance scores from figure d are plotted against the scalability scores from figure a and b. The size of the dots shows the size of the substrate.

Additionally, the moving ANCA scales better than the non-moving to unseen sizes. For example, ANCAs trained for 26 × 26 substrates comfortably scales down to 5 × 5 without noticeable performance loss. Equally, 7× 7 models scale up to 26× 26 with only a slight drop in performance. Below 7× 7, the ANCA is not scalable. In figure 5c, a network trained on 15 × 15 was scaled up to 26 × 26, and the hidden-, action- and class channels are preserved between scales. Our findings suggest that the ANCA’s strategy is retained at many scales.

Scalability offers two benefits: First, new columns with sensors can be added to the system zero shot, and perhaps future work could establish whether this can apply to new modalities and limbs (as in Huang et al. (2020)). Secondly, fault tolerance can be found through redundancy, when scaling up the ANCA from its training size.

### 1.6 Scalable ANCAs are also fault tolerant

Mature rats can adapt to having their whiskers trimmed (Erzurumlu, 2003). Similarly, a robot with distributed sensors may face sensor loss due to damage or failure in real-world applications. Assuming sensor loss can be identified, we can turn off processing of that sensor. Then, we investigate whether the redundancy in the system indeed leads to zero-shot resilience to damage, comparing the ANCA, the non-moving ANCA, a CNN, and a pre-trained Vision Transformer (ViT) from Dosovitskiy et al. (2020).

Sensor loss was simulated by setting select positions in the substrate to 0. The analogue for CNNs was chosen to be setting select positions of the first feature map to 0. Similarly, select patches in the ViT were silenced after the initial embedding layer. Loss patterns were cohesive (rectangular regions) or random and held constant throughout an episode.

We establish that the ANCA is indeed fault tolerant to sensor loss zero-shot (figure 5d). In fact, up until 70% loss, performance is maintained in all but one of the ANCAs. A similar trend is observed for random silencing, although performance declines for more classes (section B). Again, the ANCA has the distinct benefit of movement: As the substrate is silenced, the full image can still be viewed by moving the fields. In contrast, silencing leads to information loss for the CNN, ViT, and the non-moving ANCA. A moving kernel can assume a silenced kernel’s role; a non-moving kernel cannot. Next, we explore how fault tolerance relates to scalability. Fault tolerance was quantified as the performance retained with 0-90% sensor loss; scalability, as performance retained across substrate sizes (see section 3.7.2). Figure 5e shows a linear relationship for the moving ANCA: Networks excelling in scalability also excel in fault tolerance, while low-scoring networks are weak in both metrics. For the non-moving ANCA, the relationship seems weaker.

## 2 Discussion

All animals perceive while moving and through movement - as will be the case for embodied robots. Considering movement’s ubiquitous role in perception, and the mounting evidence that sensory and motor cortices collaborate, one can conclude that perception and movement are intrinsically linked. Yet, despite this emerging understanding from neuroscience, AI models inspired by the neocortex do not tend to incorporate movement.

In this work, we introduced the Active Neural Cellular Automata, a neocortex inspired system that incorporate sensor movement. We find that the system learns to classify traditional machine learning benchmarks. Importantly, we document the emergence of active sensing in the ANCA, where the kernels learn to use their movement to purposefully explore images. In fact, we observe that the ANCA displays attentive behavior, in that the fields congregate on salient points and actively follow paths that are informed by their beliefs. As opposed to a distributed system with no active sensing, the ANCA has such benefits as scalability, fault tolerance, and simplifying classification problems.

Theoretically, there are more advantages to distributed architectures. For example, we often associate a larger cortical sheet with larger intelligence. However, while ANCAs and similar architectures are highly scalable, and may lead to increased classification speed (Hawkins et al., 2017; Mousavi et al., 2019), there is no accuracy gains with larger substrates. This limitation may stem from how consensus is reached among columns. Hawkins et al. theorize that columns reach consensus through voting (Hawkins, 2021, p. 99) (as in belief averaging Galton (1907)). However, averaging beliefs dilute higher quality beliefs. Instead, consensus through local information exchange over time often yields better results in complex problems (Krause et al., 2010; Sumpter and Pratt, 2009). Although the ANCA employs local communication, its scalability leads to homogeneous behavior, narrowing the distribution of beliefs. Perhaps the key lies in incorporating the inherent architectural heterogeneousness found in the neocortex (Meyer et al., 2013), which would widen the distribution of beliefs. Then, we might indeed be able to increase performance few-shot, simply by expanding the substrate.

Further work with the ANCA requires more efficient training, as training time in evolutionary computing can be prohibitive. Considering similar approaches with distributed, message passing ensembles (Pathak et al., 2019; Huang et al., 2020), deep reinforcement learning is a powerful alternative to making further studies more feasible. Moreover, the ANCA can be used in dynamic tasks where agent action is required every timestep, for example tracking objects or controlling bodies.

Overall, our findings show that including movement in distributed systems gives rise to attentive behavior, which itself leads to many advantages. While the neocortex’s optimality suffers under evolutionary constraints that may not align with the needs of artificial systems, our results suggest that drawing inspiration from its involvement in active sensing leads to valuable emergent behaviors. With this, our work joins both the emerging body of work on neuroscience-inspired AI and the literature on the capabilities of distributed systems. Whether this architecture represents an evolutionary artifact or a scalable and resource-efficient alternative to monolithic systems remains an open question. However, its unique strengths point to a promising potential for advancing the design of intelligent systems.

## 3 Methods

### 3.1 The architecture

The ANCA has one internal feed forward network that is optimized and used as a convolutional kernel for the image and substrate. The input to the kernel is its substrate neighborhood (*K* × *K*) and its receptive field (*R* × *R*). It then outputs a substrate update, which is added to the substrate, as well as an action output *a*_*x*_ and *a*_*y*_.

The action outputs change the positions of the receptive fields. This is done by taking the raw action values *a*_*x*_ and *a*_*y*_ and thresholding them to −1, 0, or 1. This is defined for an action *a* as

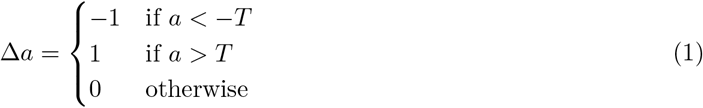

where Δ*a* is the action that will be used to displace the receptive field. The threshold value of T = 0.0007 was chosen after initial tuning to encourage the system to move early on in optimization. The output movement value *p* is then added to the current sensor position.

In addition, each column can receive its current receptive field position as input. The current position will be a normalized value of its position in the image, so that any image coordinate *x*_*image*_ and *y*_*image*_, denoted as *p*, is normalized into *p*_*normalized*_ as such:

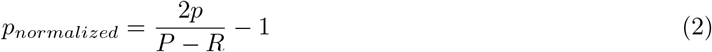

where *P* is the corresponding image dimension *N* or *M*, and *R* is the size of the kernel’s receptive field. Because the (*x*_*image*_, *y*_*image*_)-coordinate refers to the top left corner of the receptive field, *R* is subtracted from P so that the rightmost coordinate *P* − *R* will get the position 1. Note that *p*_*normalized*_ will be discretized (f. ex. with P=5 the possible values of p are −1, 0, and 1), positions are not continuous as kernels must occupy a whole pixel. This also leads to the position 0 not always being present, as it would be between 2 pixels (f. ex. for P=6, possible p is −1, −0.33, 0.33, and 1). See figure 7b for an example of what positions are received as input at the first timestep.

The initial position of each field at the start of each episode is given by stretching/squeezing the column grid (of shape *N*_*neo*_ × *M*_*neo*_) to evenly cover the image. In practic[e, this is a nea]rest neighbor mapping from the substrate positions [*x*_*neo*_, *y*_*neo*]_ to the image positions [*x*_*image*_, *y*_*image*_]:

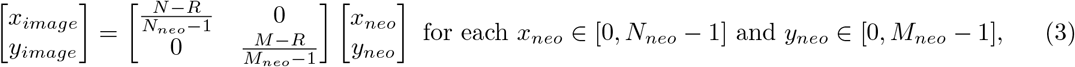

where 1 is subtracted because the code is implemented with 0-indexing. The substrate should map to a position where the full kernel will get input from the image, and so *R* is subtracted from *N* and *M*. The array [*x*_*image*_, *y*_*image*_]is rounded to the nearest integer. See figure 7a for an example mapping.

The feed forward network is intended to be as simple as possible (see Figure 6). It has one hidden layer, with a tunable number of hidden neurons, and an output layer. There are only linear activations, except in the CIFAR-10 network, which uses ReLU for all but the final layer.

**Figure 6:**
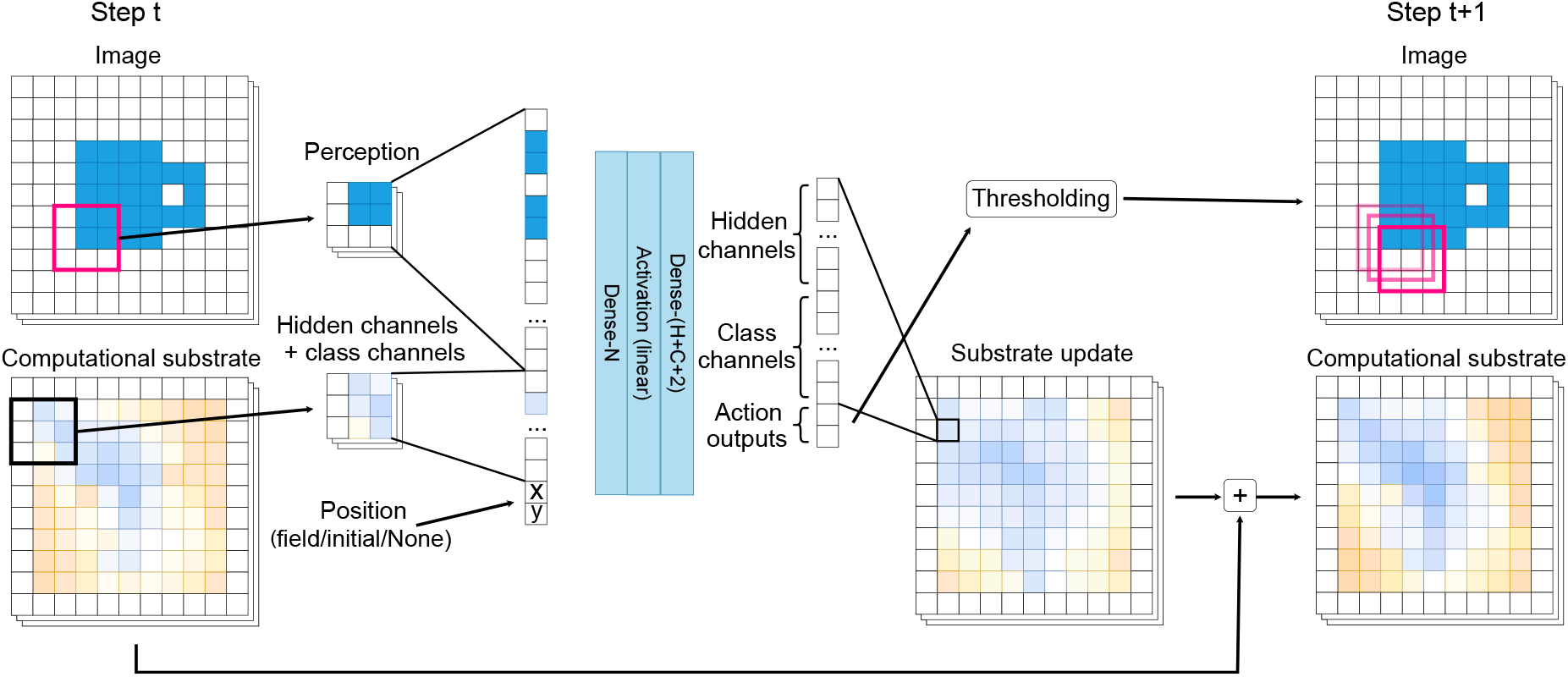
The ANCA architecture. At timestep *t*, a kernel’s receptive field (perception) and substrate neighborhood (hidden and class channels) are concatenated into one large vector. The kernel’s substrate position is alternatively appended. This input vector is processed by the network, which outputs a substrate update. The output vector has size H+C+2, for the H hidden channels, the C class channels, and the 2 actions. The actions are thresholded. At timestep *t* + 1, the substrate update has been added to the substrate and the receptive field has been moved according to the action.

**Figure 7:**
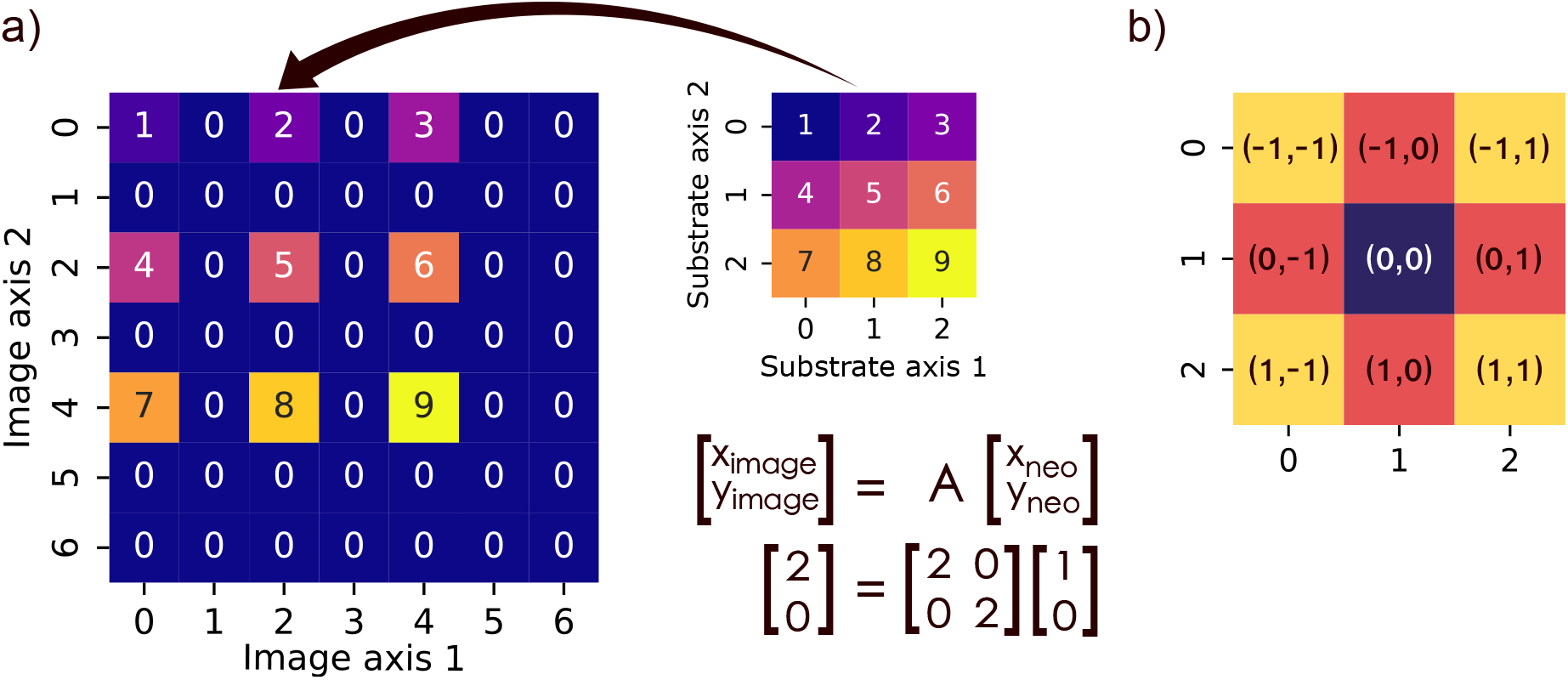
The mapping of substrate networks to positions in the image. a) Here, network 2 is shown to be mapped from substrate position [*x*_*neo*_, *y*_*neo*_] = [1, 0] to image position [*x*_*image*_, *y*_*image*_] = [2, 0]. b) The positions received by each network in their positions in a). For example, network 2 is in the topmost row and middle column, and therefore receives (−1, 0). Possible positions for a) are −1, −0.5, 0, 0.5, and 1. When network 2 moves to the left, it will receive (−1, −0.5) as input position.

The input size to a kernel in the ANCA is in general

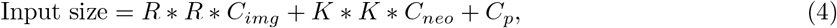

where *R*×*R* is the size of the receptive field, *C*_*img*_ is the number of channels in the input image (usually 1 or 3), *K* × *K* is the size of the substrate neighborhood the kernel takes in, and *C*_*neo*_ = *H* + *C* is the amount of hidden and class channels. Position coordinates *C*_*p*_ = 2 is added when position information is included. If not included, *C*_*p*_ = 0.

Output size is in general

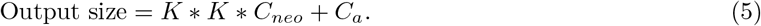

Here, action outputs *C*_*a*_ = 2 are included to specify the kernel’s action for the receptive field. When action is not included, *C*_*a*_ = 0.

### 3.2 Training

The system is trained using Covariance Matrix Adaptation-Evolutionary Strategy (Hansen and Ostermeier, 1996). The first solutions for the network weights are generated by CMA-ES using a multi-variate Gaussian distribution with mean equals to 0 and standard deviation equals to 0.002. Using multiprocessing, the optimization evaluates the population across as many threads as there are individuals in the population. The optimization is run until the maximum number of generations is reached.

We train and test our model using three common datasets for image classification, MNIST (Deng, 2012), Fashion-MNIST (Xiao et al., 2017), and CIFAR-10 (Krizhevsky et al., 2009). All of them uses a ca. 85/15 train-test-split. During training, a small batch is randomly selected to record the networks’ losses. Every 200 generations, the network with the lowest loss gets tested on another training batch to record its performance on data not seen in this generation. The test data is not used during training, but rather during testing when considering performance, behavior, scalability, and fault tolerance. Finally, the most optimized network is used in all tests.

### 3.3 Loss and accuracy

The prediction operator is defined as

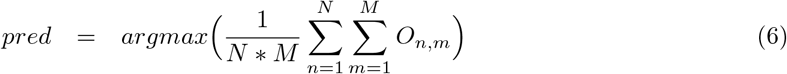

where N × M is the class channel dimensions, and *O*_*n,m*_ denotes one pixel of size C in the class channels.

The predicted value is used in the accuracy:

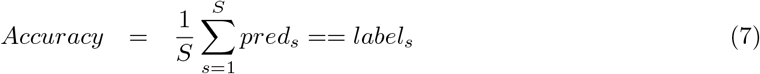

where S is the number of image samples, and *label*_*s*_ is the corresponding correct label to sample *s. pred*_*s*_ is the system’s prediction on the image, as defined in equation 6. The accuracy is not a continuous value and therefore is not used to optimize the system. Rather, the accuracy helps determine if the system solves the classification task.

Through initial trials, we discovered that Categorical Cross Entropy (CE) gave better accuracy than Mean Squared Error (MSE) when optimizing the system. However, while MSE keeps the system output values around 1 and 0, CE makes the output values explode as the system evolves through the episode. Therefore, CE was paired with minimization of the output class channel values:

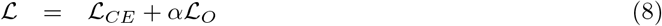

Where ℒ_*CE*_ is the CE loss and ℒ_*O*_ is the sum of the output class channel values. *α* defines a scaling factor of the regularization term, defined below. Both loss terms are defined for each pixel n,m in the output channels of size *N*_*neo*_ × *M*_*neo*_:

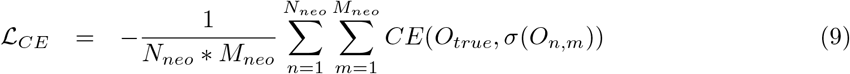

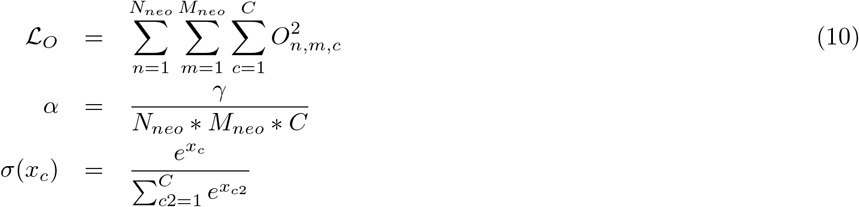

where *C* is the number of output class channels (e.g. for 5 classes, C=5), *O*_*true*_ is the true sample label (one-hot encoded), and *O*_*n,m*_ is the pixel vector of size C indicating the kernel’s classification confidence. *α* was introduced to scale the output channel minimization with the number of classes and *N*_*neo*_ and *M*_*neo*_ throughout the experiments. *γ* is the normal scaling factor of the regularization term. It was set to the value 1125*λ*. Because the system was primarily developed while testing on 5 classes with a 15 × 15 computational substrate, the number 15∗ 15 ∗ 5 = 1125 was used as scaling so that any other combination of number of classes and size of substrate would not make the regularization more or less severe. This way, the system could scale up without increasing the regularization loss. *λ* was set to 10^*−*4^.

In addition, for CIFAR-10, weight regularization was added to the loss instead of regularization of output values. This was defined as

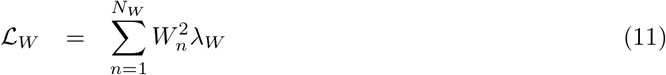

for the *N*_*W*_ flat weights *W* (all weights and biases, flattened). *λ*_*W*_ was set to 10^*−*4^.

### 3.4 Hyperparameter choices

As can be seen in table 2, many hyperparameters have to be tuned for each ANCA. For the training with CMA-ES, the general tuning philosophy was that more is better. However, being limited by hardware, we have chosen to run the optimization for approximately 50k-120k generations and 100 in population size. The initial standard deviation, *σ*, is set to 0.001-0.005, but has not been tuned extensively. Again, for samples per class, more is better, but again due to hardware constraints, 20 to 50 was chosen depending on the dataset. The regularization factors *λ* = 0.0001 and *λ*_*W*_ = 0.0001 were chosen as small values to scale the penalizations.

**Table 2:**
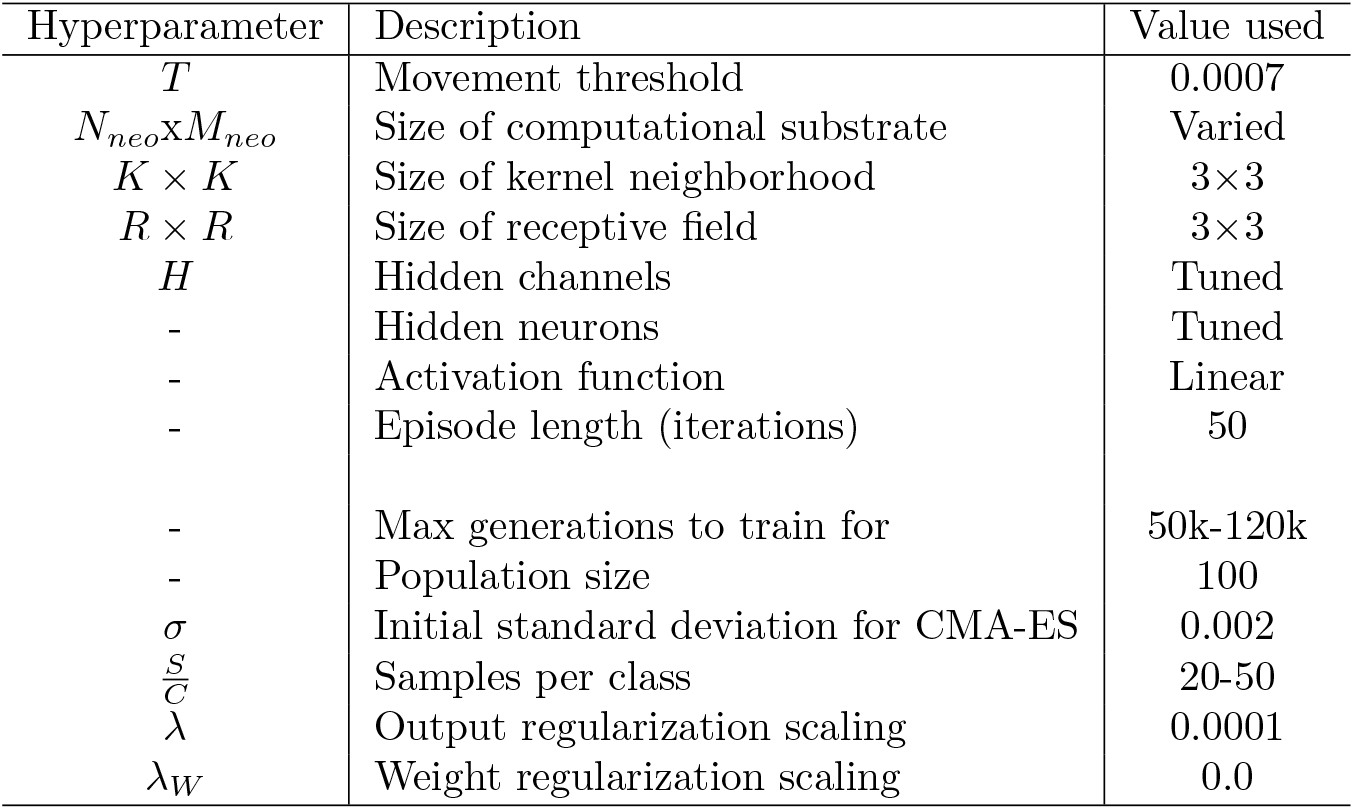
The default hyperparameters of the system. The upper hyperparameters define the system, while the lower ones define the training as required by CMA-ES. As opposed to a naive NCA, this method adds three new parameters: *N*_*neo*_ × *M*_*neo*_, *T*, and *R* × *R*.

The movement threshold *T* = 0.0007 was chosen to encourage movement early in the optimization. Because the network is zero-initialized, and CMA-ES uses a small *σ* to update the weights, the initial outputs of the network were quite small. The chosen value *T* = 0.0007 allowed the network to move a little after a few generations, without always moving to a corner. Because CMA-ES is more exploitative than exploratory, especially with a low *σ*, this helped make the optimization faster. This value was found by trial and error and may not be optimal.

The size of the computational substrate was chosen to be 7×7 (for most of the datasets) because it is faster to train than the image size (28×28). The number of iterations for the episodes were tested for 5 classes of MNIST, and it seemed as a rule that more iterations were better. However, around 50 iterations, the performance gain from more iterations started to become vanishingly small, and so 50 iterations were chosen.

The only hyperparameters that were actively tuned for each dataset were the number of hidden channels and hidden neurons. As they determine the number of parameters, they also determine both the complexity of the model and the speed of the optimization. The latter of which became important as CMA-ES converges faster, and is less prone to failure in convergence, with fewer parameters (Müller and Glasmachers, 2018). Therefore, for the image datasets, a manual grid search was conducted on the hidden channels (1 to 10) and the hidden neurons (10 * (1 to 9)). Not every grid was tried if a promising area was found by hand. The final hyperparameters can be seen in table 3, and more extensively in the Appendix (Section F).

**Table 3:** The final hyperparameters chosen for the different datasets for table 1 and their training time.

| Dataset | Hidden channels | Hidden neurons | Total parameters | Substrate size | Training time |
| --- | --- | --- | --- | --- | --- |
| MNIST 5 classes | 5 | 20 | 2292 | $7 \times 7$ | 1 day |
| MNIST 10 classes | 5 | 40 | 6577 | $7 \times 7$ | 7 days |
| Fashion-MNIST | 5 | 40 | 6577 | $7 \times 7$ | 9 days |
| Fashion-MNIST | 5 | 40 | 6577 | $15 \times 15$ | 16 days |
| CIFAR-10 4 classes | 2 | (20, 20) | 2268 | $15 \times 15$ | 7 days |

For the cup dataset, we chose to use many hyperparameter pairs to explore how the results held for different tunings. Therefore, combinations of hidden channels (1-3) and hidden neurons (10, 20, 30, 40) were trained four times, resulting in 48 networks per condition. The results found in section 1.4 holds for all hyperparameter pairs used. More hidden channels and neurons was not seen as necessary because the task is so simple.

### 3.5 The data

For this project, 4 datasets were used: One generated toy-dataset, and MNIST, Fashion-MNIST, and CIFAR-10. In all cases, the labels were one-hot encoded.

The toy-dataset were the 3-class simple objects (the cup-data). Because the data was generated, it only had pixel values 0.0 and 1.0. Every generation was trained on the 3 images, as there was no variation in the data, and no test set.

The image data is normalized to the interval [0, 1]. Every generation, a balanced set is randomly chosen from the full datasets corresponding to the number of samples per class. For example, for 10 samples per class, and 5 classes, a balanced batch of size 50 is chosen for each generation. This is done to avoid training on batches without all the classes, and to give equal importance to each class.

### 3.6 Displaying ANCA behavior

The ANCA moves its fields every timestep, with corresponding changes to the substrate. Three methods were chosen to display its behavior.

The first is to simply display the fields as 3 × 3 squares at the field position in the image. This is the method chosen for the videos and figure 4b. However, this does not show if fields are overlapping, and what beliefs they have in those positions.

The next method is used only for displaying focusing behavior (figure 2). Here, a matrix is collected for every timestep with Boolean values: If the pixel was visited or not by a field. Then, these matrices are averaged for a final image. These figures give an idea of where fields spend their time, but do not include how many fields congregate at one coordinate within a timestep. This was purposefully left out because ANCAs with high internal memory (many hidden channels) choose to go towards a corner at the end of an episode. This could make it difficult to see what they spent their time on during the episode (without purposefully leaving out timesteps).

Lastly, we sought to make a plot similar to a vector field to show what positions led to which actions (figure 3). To make the actions clearer, we averaged actions across 2 timesteps. However, as a dynamical system, the behavior of the ANCA depends on more than the coordinates (*x, y*): Two kernels with the same field position also differs in the substrate channels. Therefore, we chose instead to average the behavior of every kernel at position (*x, y*) over 10 timesteps and display their average action in that position as an arrow. The arrow is opaquer if more kernels contributed to it. The arrow is also colored with the current highest output to the class channels for that position. This way, the number of fields in a position, their beliefs, and their actions are displayed.

### 3.7 Ablation study details

In section 1.4, 1.5, and 1.6, versions of the ANCA were used where key features were left out. The details of how this was achieved are recounted here. In general, features that were removed were not zero-padded, inputs and outputs were simply removed, which only slightly reduces the number of parameters in the system.

To exclude position information, the current position was not added to the input for each kernel. This removes two input neurons. Likewise, for excluding movement capabilities, output neurons are simply reduced, and receptive field positions remain as initialized. Output size is then also reduced by two.

When movement is not included, the ANCA becomes a naive NCA. However, this kind of naive NCA has not been used for classification at the time of this paper. Randazzo et al. (2020) and Walker et al. (2022) reduce the irrelevant information that goes to the NCA by only processing kernels that overlap the object. As such, the ablation study is not meant to be seen as a comparison to NCAs, but simply to study the effect of not including movement and position when all other features are kept constant.

#### 3.7.1 Fault tolerance and scalability

Further details about the experiments in section 1.5 and 1.6 are presented here.

For figure 5a and b, networks were trained on substrate sizes *x × x* where *x* was 1, 3, 5, 7, 15, and 26. For all sizes, 3 networks were collected. 40 image samples per class were used to test the performance at every substrate size.

For fault tolerance experiments, figure 5d, the networks trained with 26 × 26 substrates were reused from figure 5a and b. In addition, 5 more networks were trained for the moving and non-moving ANCA. In total, the moving and non-moving ANCA, as well as the CNN and ViT, had 8 trained networks for figure 5d.

For Figure 5e, every substrate size, *x × x* with *x* from 4 to 26, was included. Here, networks were reused from the previous experiments and a few more were trained so that, in total, 46 networks were included for each the moving and non-moving ANCA. Fewer networks were collected for sizes *>* 13 because of the increased training time.

When testing fault tolerance, 40 image samples per class were used, and a randomly placed area in the substrate (ANCAs) or feature map (CNN) was put to 0. For the ViT, the patch embeddings were put to 0, after position encoding was added. The [CLS] token was never set to 0. If the silenced area was cohesive, it was also randomly shaped in a roughly square shape, as long as it had enough pixels for the current test-size. Test-sizes were determined as percentage of the substrate/feature map/patch embeddings.

**The CNN** The CNN used for fault tolerance was implemented using Tensorflow (Abadi et al., 2015). It was tuned insofar that it guaranteed good performance. Its architecture is as follows:

1. A Convolutional layer, 10 3 × 3 kernels, with no padding and ReLU activation
2. A Max Pooling layer, a 2 × 2 kernel,
3. A Convolutional layer, 10 3 × 3 kernels, with no padding and ReLU activation
4. A Max Pooling layer, a 2 × 2 kernel,
5. Flatten, then a Dense layer, 100 hidden neurons, ReLU activation
6. Output layer of 3 neurons, and softmax activation.

It was trained for 30 epochs, with a batch size of 200. The Adam optimizer was used, with a learning rate of 10^*−*4^. As with the ANCA, the CNN was optimized with CE loss and minimization of output loss

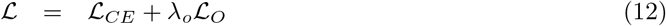

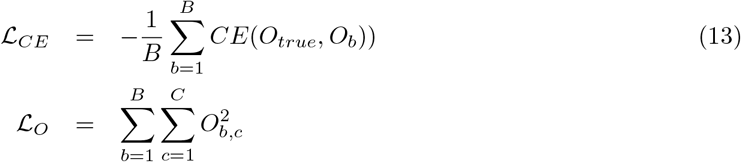

where *B* = 200 is the batch size, *O*_*true*_ is the true label, *O*_*b*_ is the corresponding predicted value, and *C* is the number of classes, which is 3 in section 1.6 and also 5 in the appendix section B. *λ*_*o*_ = 10^*−*4^ scales the penalization. Output minimization was included to keep the conditions similar for the ANCA and CNN.

Accuracy was defined as

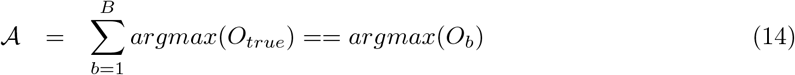

The accuracy for 3 classes on the train set was 99.93% and 99.75% on the test set. For 5 classes, it was 99.94% on the train set and 99.76% on the test set.

**The ViT** The ViT used for comparing fault tolerance was implemented using PyTorch (Paszke et al., 2017) and especially the pre-trained model of Dosovitskiy et al. (2020) through the library Transformers (Wolf et al., 2019). It has 86.4 million parameters, and was pre-trained on ImageNet-21k (Deng et al., 2009). Additional parameters were added with the classification head. Because it has been trained for RGB images of 224×224 pixels, our datasets were scaled up and converted to RGB (while maintaining grayscale). Bilinear interpolation was used. Additionally, the recommended mean and standard deviation of 0.5 per channel was used to normalize the images.

Its architecture consists of a patch embedding layer and a vision transformer encoder with 12 attention blocks, each with 12 attention heads. After patch embedding, a [CLS] token is concatenated to the list of embedded patches to account for global image information. Further architecture details can be found in Dosovitskiy et al. (2020).

The ViT was fine-tuned for max 3 epochs, with a batch size of 32. Batches were not balanced in terms of classes. We observed fast convergence with the pre-trained model, and chose to therefore have early stopping to avoid overfitting, which every 20 batch tested the validation accuracy. The validation set was 2% of the train dataset. The early stopping had a patience of 7, and a minimum difference between previous and current loss of 0.005. The learning rate was 10^*−*4^, and the model also had a small weight decay to avoid overfitting, where the weight decay parameter was 10^*−*5^.

The loss and accuracy was defined respectively as in equations 13 and 14. Hyperparameters were tuned only insofar as to give good performance. The best recorded ViT for 3 classes of MNIST got a train accuracy of 99.87% and test accuracy of 99.81%. For 5 classes of MNIST, the train accuracy is 99.80% and the test accuracy is 99.81%.

#### 3.7.2 Fault tolerance and scalability metrics

In section 1.6, metrics were needed to show the relationship between scalability and fault tolerance.

Fault tolerance is here formally defined as

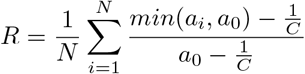

for measured accuracies *a*_*i*_ and classes *C* for measurements *N*. *a*_0_ is the performance of the system on no loss of sensors. *min*(*a*_*i*_, *a*_0_) was included to ensure no fault tolerance could be more than 100%, in the instance the model performed better with some loss of sensors. Scalability was similarly defined as

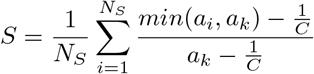

where *a*_*k*_ is the accuracy recorded for the substrate size the network was trained for, and *N*_*S*_ is the tested substrate sizes from 1 to *N*_*S*_ × *N*_*S*_. Here, *N*_*S*_=26.

### 3.8 Language model use

The Large Language Model (LLM) ChatGPT was utilized for the code and text during this project. For the code, ChatGPT was used to optimize code snippets for speed, often from pure Python to NumPy vectorization. Any code optimized with ChatGPT has been extensively tested to ensure its correctness. In addition, ChatGPT was utilized for brainstorming reasons for issues with the code. For the text, ChatGPT was used for brainstorming paragraph structure and for suggesting wording that would keep the text professional and concise. LLMs have not generated substantial parts of the code or text in this article.

## Supporting information

Supplementary video summary

## 4 Acknowledgments

The research presented in this paper has benefited from the Experimental Infrastructure for Exploration of Exascale Computing (eX3), which is financially supported by the Research Council of Norway under contract 270053. We also want to thank Trym Lindell and Stefano Nichele for their valuable input on the project.

## Appendix

### A Further active sensing

**Figure A1:**
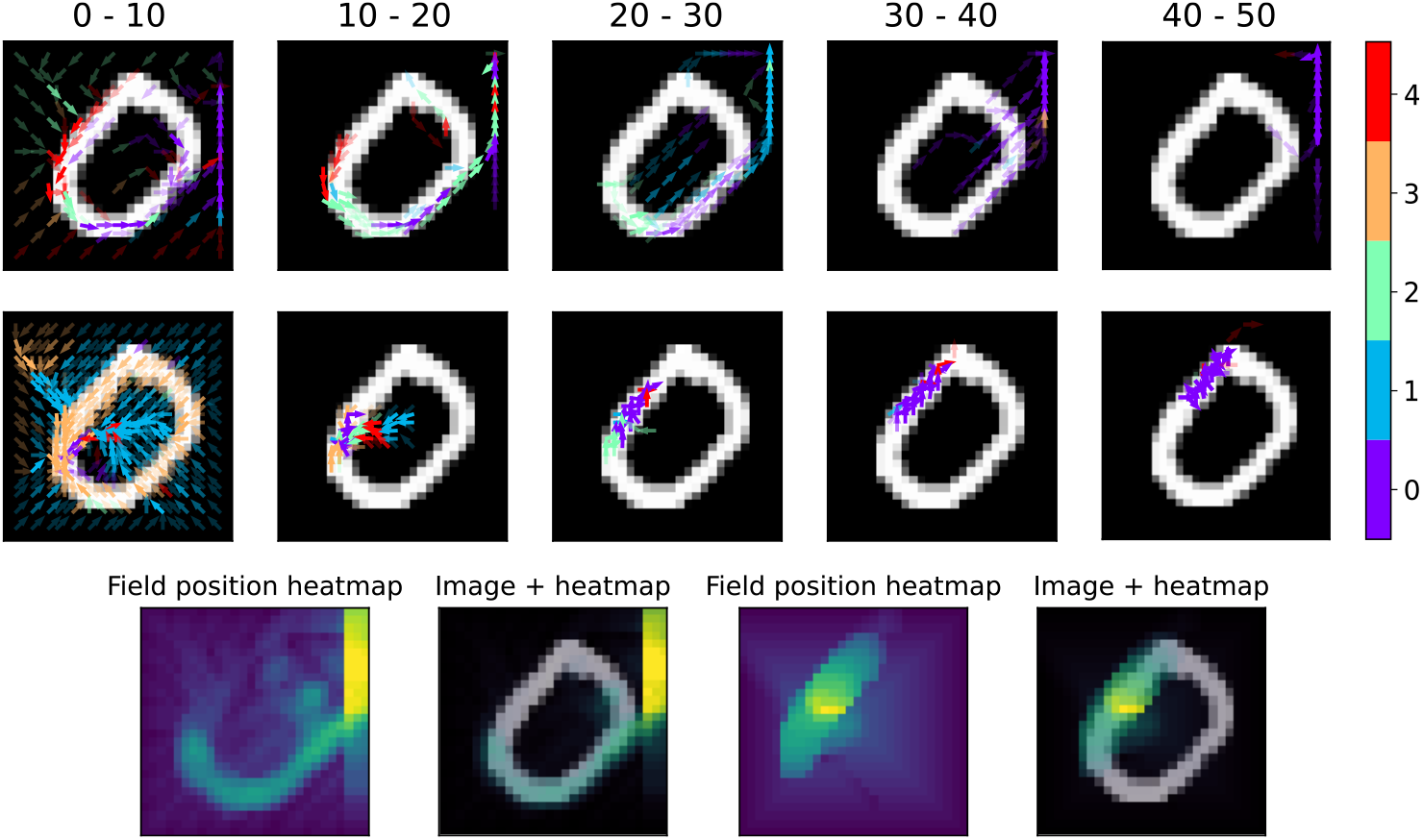
Effect of parameter choices on behavior. Top row: The behavior of the top-performer (accuracy 96.34%) with a 7×7 substrate and 5 hidden channels. Middle row: The behavior of the top-performer (accuracy 96.20%) with a 15×15 substrate and 2 hidden channels. Bottom row: Left: Focus of 7×7 substrate. Right: Focus of 15×15 substrate.

### B Further fault tolerance experiments: Fault tolerance decreases with more classes

As shown in section 1.6, the ANCA’s scalability leads to increased fault tolerance. In this section, we supplement these results with experiments on when the damage is randomly distributed instead of cohesively (in a rectangle). We also show the fault tolerance for the ANCA, the CNN, and the ViT for 5 classes.

At 3 classes, we see that the ANCA is still quite fault tolerant with random damage (Figure B2c). This type of noise hinders the information flow in the computational substrate, and so we can infer that these networks are not relying on communications to make their predictions. However, when testing fault tolerance on networks trained on 5 classes (Figure B2d), it becomes immediately clear that hindering information flow makes the ANCA’s accuracy drop. Qualitatively, the ANCA is in general not very fault tolerant against randomly applied damage, because it is reliant on information flow in the substrate.

For cohesive damage, the ANCA’s information flow is maintained, and so it can to a greater degree maintain performance. In Figure B2b, we show a vast majority of the networks trained for 5 classes still are fault tolerant towards 50% removed sensors. In this figure, the ANCA is trained with 15 × 15 substrate to facilitate scalability and therefore fault tolerance. We train 8 networks with 2 hidden channels and 40 hidden neurons, and 6 networks with 5 hidden channels and 20 hidden neurons. This was done because we saw that fault tolerance and scalability decreased with an increased number of hidden channels.

When comparing the ANCA to the ViT, another method that selectively attends the input space, we can see that they have different strengths: The ViT fares well with randomly applied silencing, while struggling with cohesive - and the ANCA struggles with randomly applied silencing but excels at cohesive. We can therefore hypothesize that the ANCA could benefit from an attention-driven aggregation operator, such as in (Tang and Ha, 2021), to further its fault tolerance to sensor damage.

**Figure B2:**
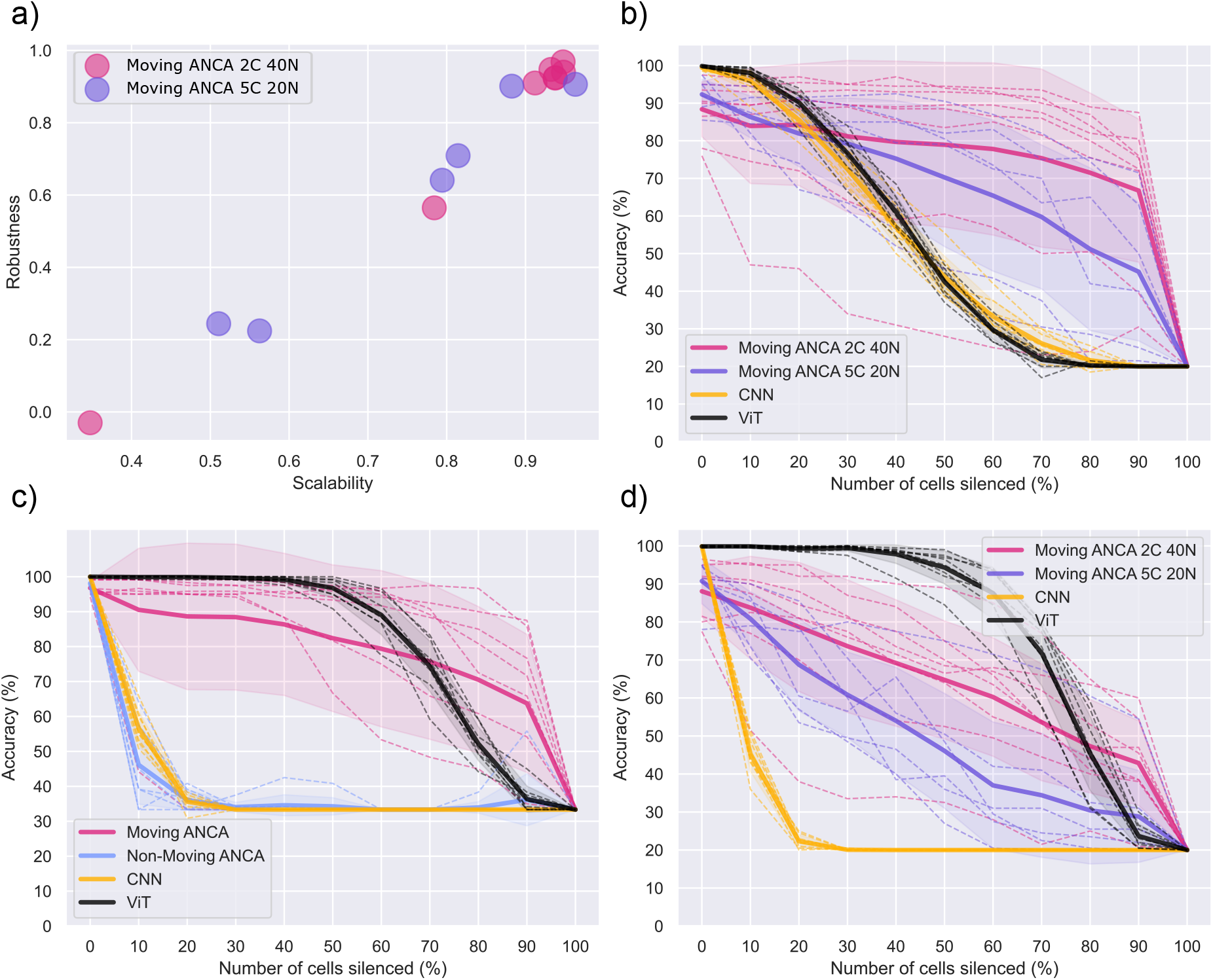
Further fault tolerance. Both Moving and Non-moving ANCAs has position information. 2C 40N refers to 2 hidden channels and 40 hidden neurons. a) The ANCA was trained for 5 classes of MNIST and was then measured for scalability and fault tolerance against sensor silencing. At 5 classes, there still is a linear relationship between these two quantities. b) The recorded fault tolerance to cohesive sensor silencing for a CNN, a ViT, and two ANCAs with different sized substrates. c) For 3 classes of MNIST, the moving and non-moving ANCA, a CNN, and a ViT are compared for randomly silenced sensors. d) As in b), but with randomly silenced sensors.

### C Further ANCA focus plots

**Figure C3:**
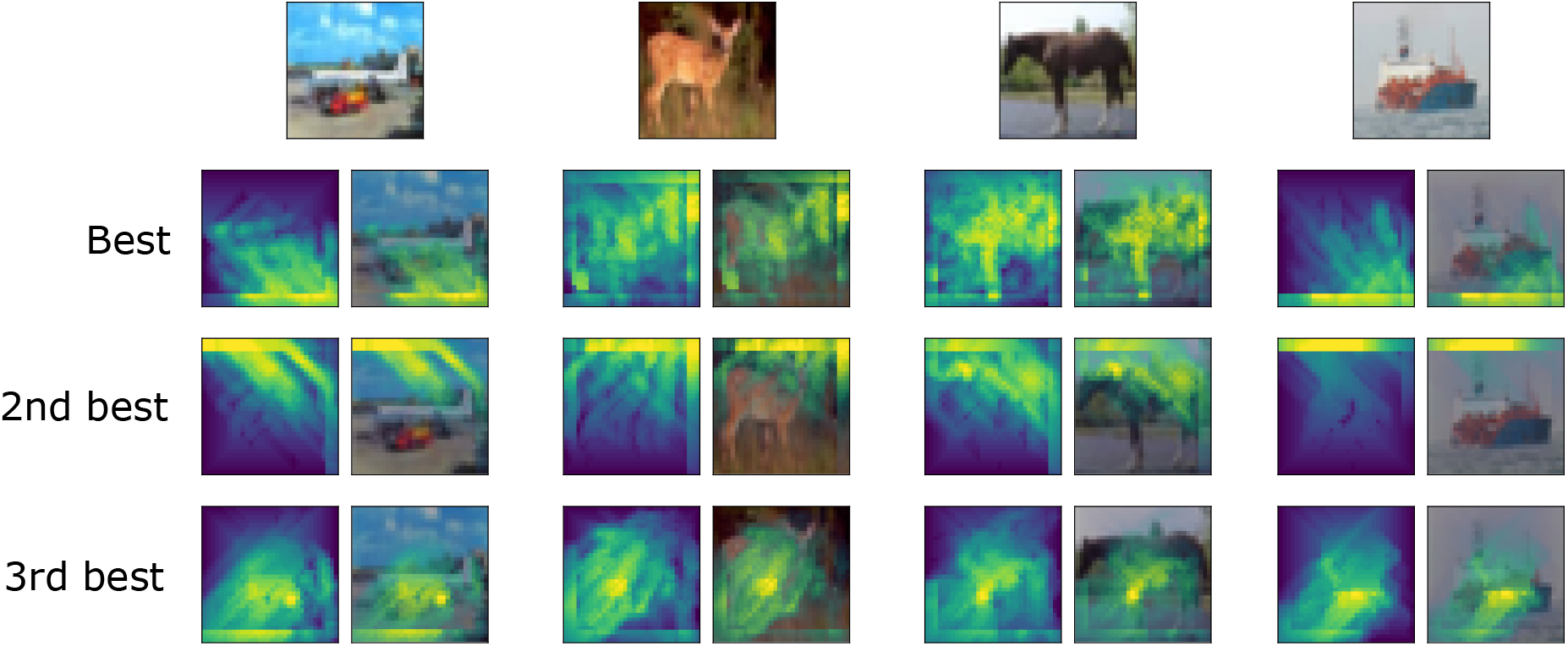
Further focus for MNIST and Fashion-MNIST. The best and second best performing ANCAs are shown. Misclassifications are marked in red.

**Figure C4:**
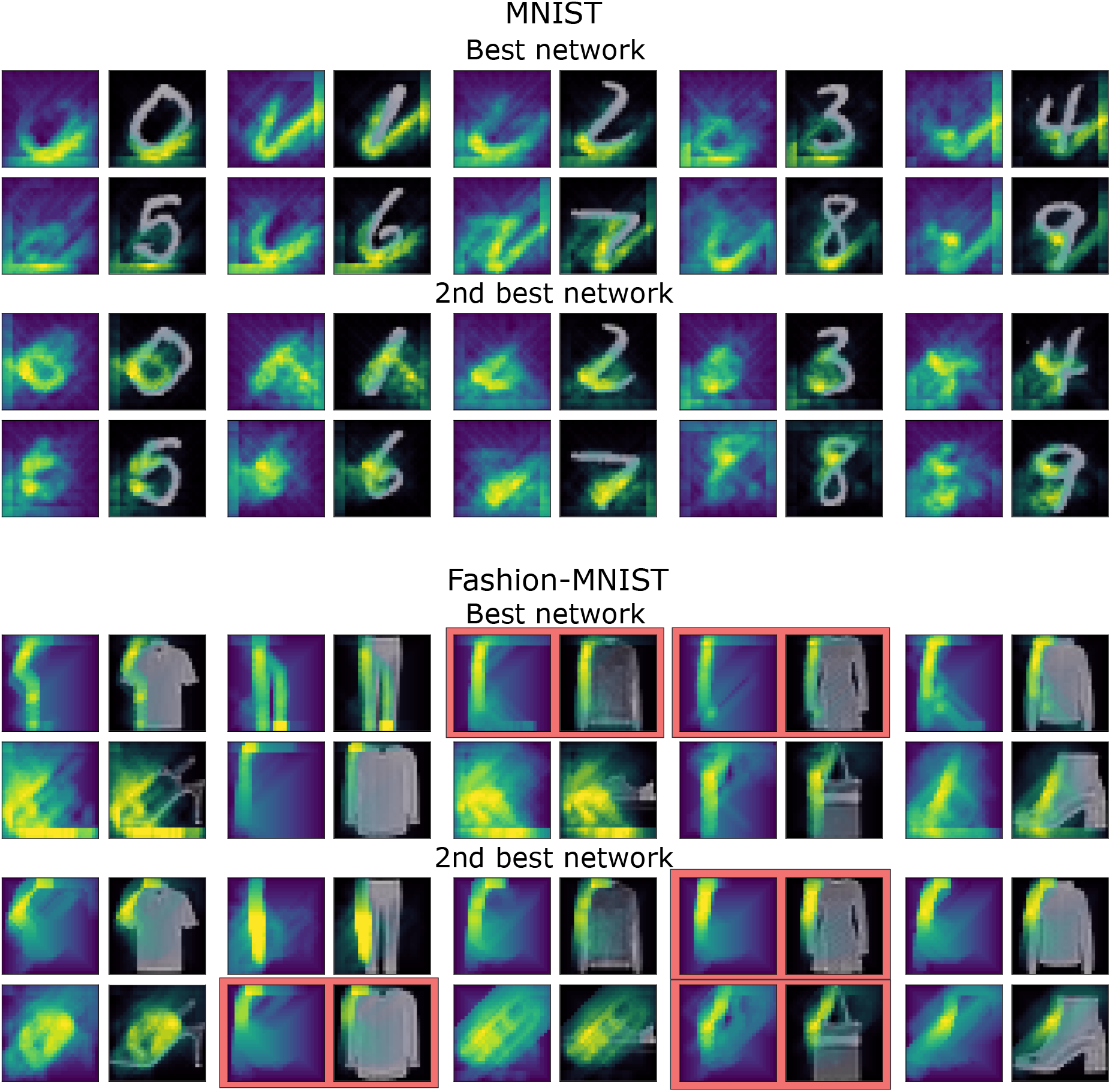
Further focus for CIFAR-10 4 classes. The best and second best performing ANCAs are shown. There are no misclassifications here. For this dataset, the networks reveal that they focus largely on the background. For example, the 2nd best network focuses exclusively on the top of the image, suggesting that it would not generalize well to the same object in a different context. Compare that to the 3rd best, which spends most of its time on the object. This feature makes the system more interpretable, and you can more easily see which network might generalize the best.

### D Complete list of accuracies

**Table 4:** The accuracies of the toy dataset. There is no test set, so only the train accuracies are reported. All methods had networks that solved the task. However, the mean accuracy was much higher for the moving systems because some non-moving systems could not classify more than 2 out of 3 classes consistently.

|  | Moving | Position | Best train | Mean train | # runs |
| --- | --- | --- | --- | --- | --- |
| Simple Object | True | True | 100.0% | 99.31±4.8% | 48 |
|  | True | False | 100.0% | 99.31±4.8% | 48 |
|  | False | True | 100.0% | 82.64±16.7% | 48 |
|  | False | False | 100.0% | 79.86±16.3% | 48 |

**Table 5:** The accuracies of benchmark datasets. Different sizes of substrates N × M were tested, as well as different number of hidden channels H.

|  | N×M | H | Best train | Best test | Mean train | Mean test | # runs |
| --- | --- | --- | --- | --- | --- | --- | --- |
| MNIST 5 class | 7x7 | 5 | 96.34% | 97.27% | 93.61±1.8% | 94.50±1.8% | 8 |
|  | 15x15 | 5 | 93.81% | 94.71% | 90.75±4.1% | 91.53±4.2% | 6 |
|  | 15x15 | 2 | 95.77% | 96.20% | 87.63±7.0% | 88.33±7.3% | 8 |
| MNIST 10 class | 7x7 |  | 91.57% | 91.97% | 88.99±1.9% | 89.53±1.7% | 7 |
| Fashion 10 class | 7x7 |  | 76.45% | 76.04% | 75.08±1.3% | 74.36±1.4% | 8 |
|  | 15x15 |  | 74.54% | 73.37% | 71.98±1.6% | 70.97±1.6% | 4 |
| CIFAR-10 4 class | 15x15 |  | 69.37% | 70.70% | 66.86±1.9% | 68.13±1.8% | 4 |

### E Confusion matrix for Fashion-MNIST and behaviors

The confusion matrix in figure E5 was sorted by ANCA grouping. That is, Torso garments (dress, t-shirt, pullover, coat and shirt) are displayed next to each other. The behavior for bag for this particular network was also similar to torso garments and is displayed afterwards. Trouser is a unique behavior and is displayed after bag, and all shoes follow. It appears that confusion is largest within groups. Especially torso garments lead to a large amount of confusion. Perhaps the ANCA would do better with larger receptive fields, to help clarify small patterns such as collars and hoods.

**Figure E5:**
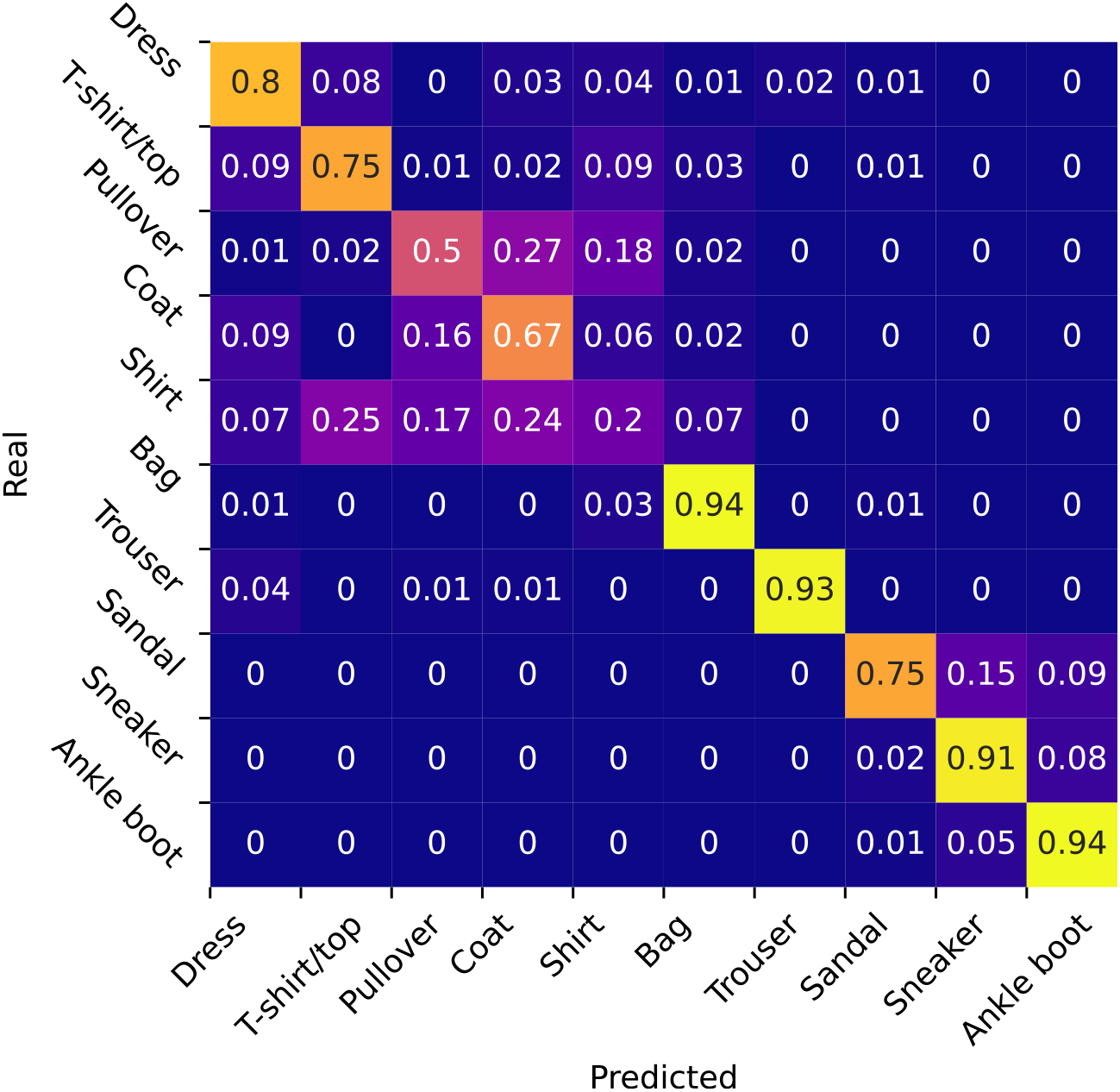
The confusion matrix of the top performing Fashion-MNIST ANCA with a large substrate (15 × 15). The confusion matrix is made of 500 samples per class, and sorted by group behavior (torso garments first, then bag, trousers, and shoes).

The distinct group behaviors for the top performer with a large substrate on Fashion-MNIST is given in figure E6. Here, the grouping is pointed out. This grouping behavior is fairly consistent for ANCAs on Fashion MNIST: Both the best and second best ANCA displays the same behavior. However, the other two shows distinct behaviors only for shoes. Additionally, the top performer on Fashion-MNIST with a small substrate also shows this exact grouping behavior.

**Figure E6:**
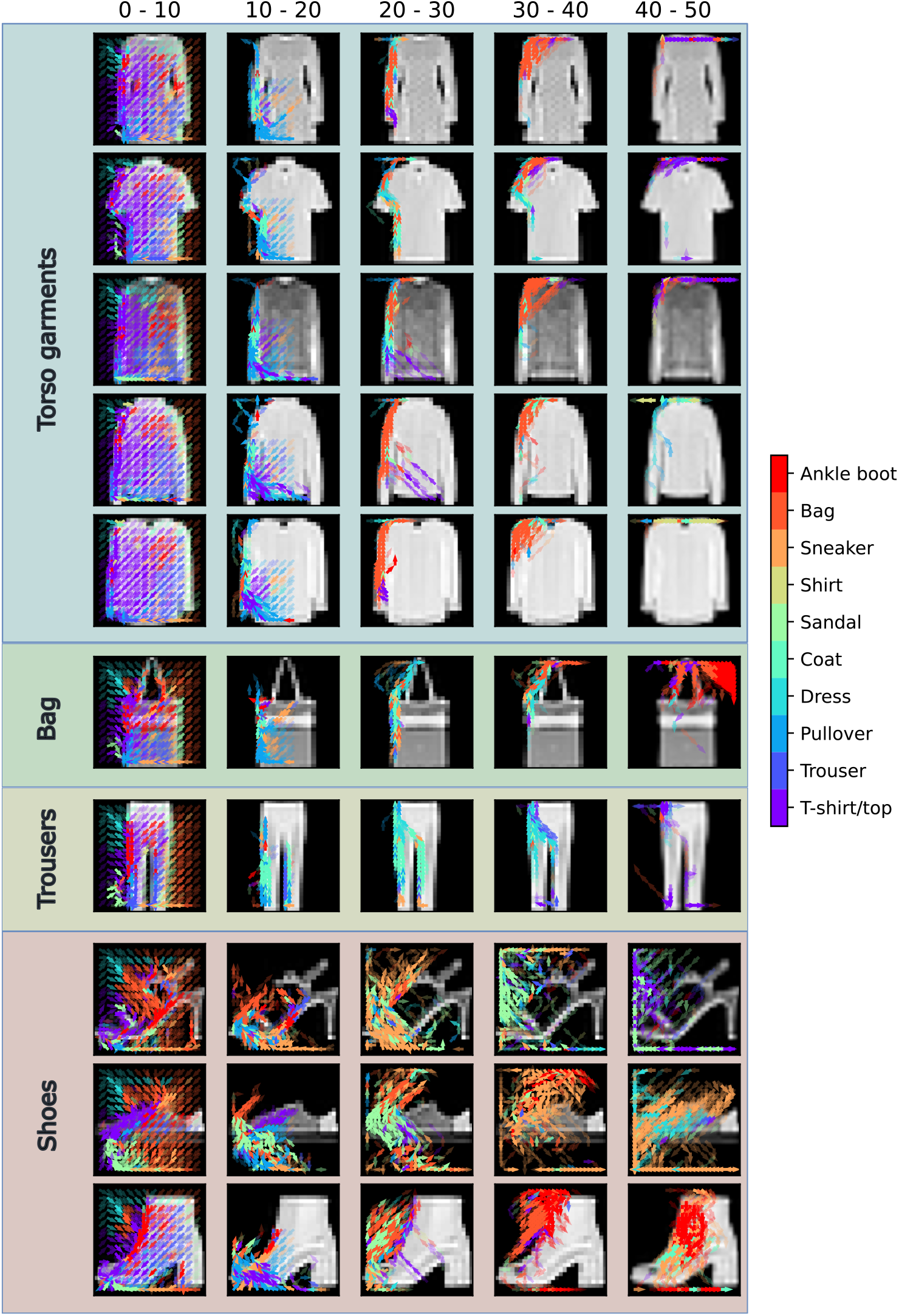
All behaviors of the top performer with a large substrate on Fashion-MNIST. The groups are marked, with bag as a possible additional group, on top of torso garments, trousers, and shoes. Among these, two classes are misclassified: Dress (classified as coat), and pullover (classified as shirt).

A similar grouping behavior can be seen for MNIST, but it is much less prominent. In figure E7, we see that there appears to be a grouping of pointy versus round digits. And although this behavior can be seen across several trained ANCAs, it is far less consistent than for Fashion-MNIST. Indeed, for larger substrates, the behavior might be completely homogeneous between digits. However, for smaller substrates, some sort of grouping is usually present along the lines seen in figure E7.

**Figure E7:**
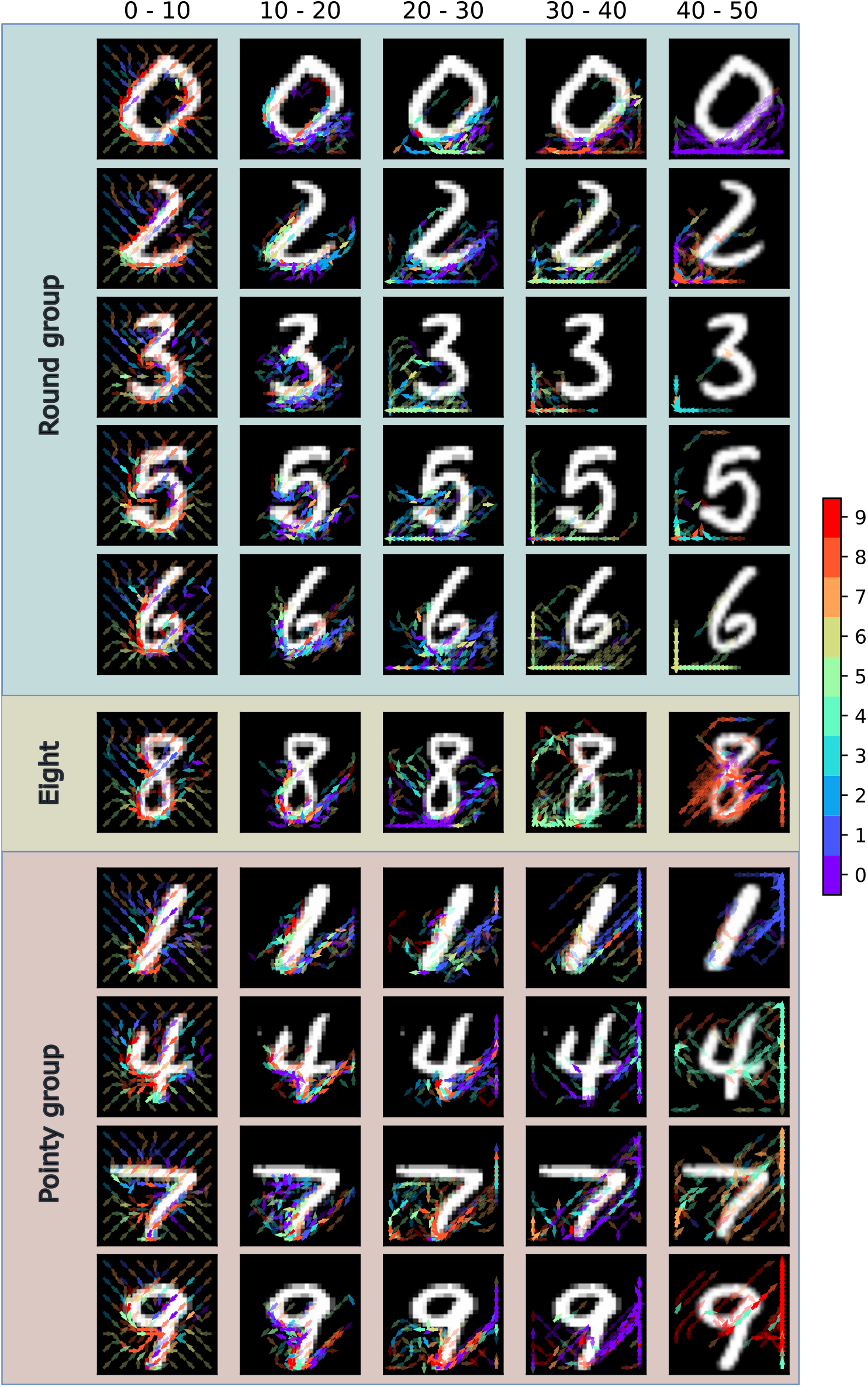
All behaviors of the top performer on MNIST. Everything is correctly classified. The initial behavior is very similar for all digits: Crowd to the edges and follow them counterclockwise. However, from timestep 20 and onward, there seems to be a distinct behavior between the groups of pointy and round digits. Note especially that towards the end of the episode, the fields retreat to a corner that is similar within groups. The behavior on digit 8 seems to be somewhat of an outlier, initially following the “round” group behavior, but then returning to the middle of the digit.

### F All hyperparameters

**Table 6:** The hyperparameters for the cup data that was used in section 1.4. Hidden channels and hidden neurons were varied between these values to see how tuning would affect the convergence. Values were kept relatively low because the task is very simple. No matter the tuning, the results in section 1.4 holds.

| Hyperparameter | Description | All setups |
| --- | --- | --- |
| $N_{neo} \times M_{neo}$ | Size of computational substrate | $16 \times 16$ |
| $H$ | Hidden channels | 1, 2, 3 |
| - | Hidden neurons | 10, 20, 30, 40 |
| - | Activation function | Linear |
| - | Max generations to train for | 20k |
| - | Population size | 30 |
| $\sigma$ | Initial standard deviation for CMA-ES | 0.001 |

**Table 7:**
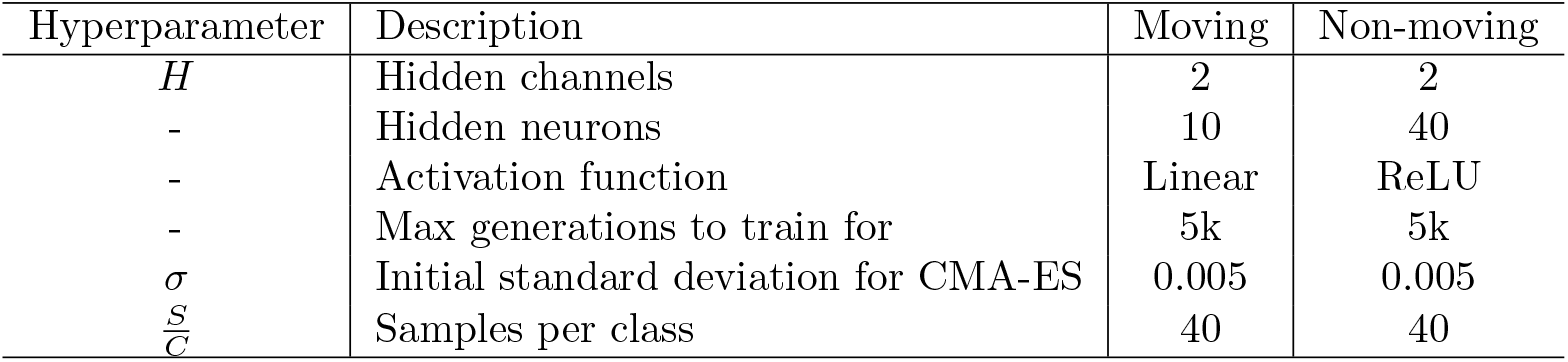
The hyperparameters for 3 class MNIST. used in section 1.5, 1.6, and B. Both setups were hand-tuned to find a good performance. The moving system was easier to tune, while the non-moving needed a bit more work, and ended up with ReLU and more hidden neurons to converge in the same time as the non-moving.

**Table 8:** The hyperparameters for 5 class MNIST. All setups were hand-tuned to find a good performance. These three systems were used in section 1.3 to compare substrate sizes.

| Hyperparameter | Description | Setup 1 | Setup 2 | Setup 3 |
| --- | --- | --- | --- | --- |
| $N_{neo} \times M_{neo}$ | Size of computational substrate | $7 \times 7$ | $15 \times 15$ | $15 \times 15$ |
| $H$ | Hidden channels | 5 | 5 | 2 |
| - | Hidden neurons | 20 | 20 | 40 |
| - | Max generations to train for | 50k | 50k | 50k |
| $\sigma$ | Initial standard deviation for CMA-ES | 0.002 | 0.002 | 0.001 |
| $\frac{S}{C}$ | Samples per class | 30 | 30 | 20 |

**Table 9:**
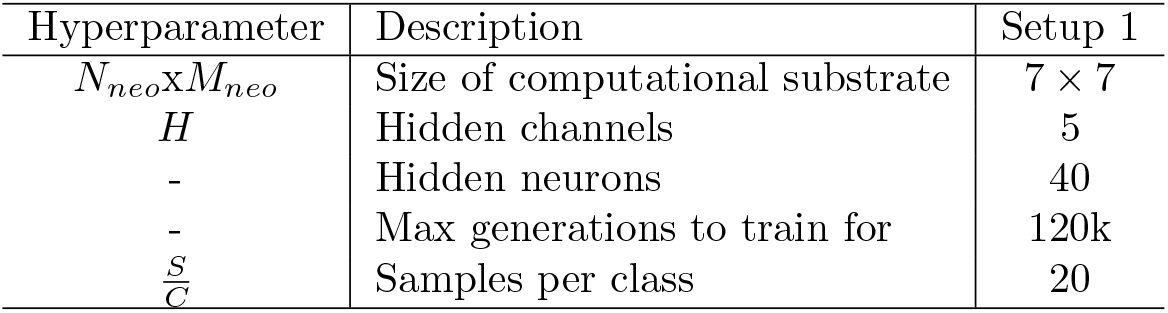
The hyperparameters for 10 class MNIST. The setup was hand-tuned for best performance.

**Table 10:** The hyperparameters for Fashion-MNIST. Setup 1 and Setup 2 had the same parameters as MNIST 10 classes, but with fewer generations because of computational constraints. Setup 1 had more samples per class because the task has more variance than MNIST, however Setup 2 could not because of training time.

| Hyperparameter | Description | Setup 1 | Setup 2 |
| --- | --- | --- | --- |
| $N_{neo} \times M_{neo}$ | Size of computational substrate | $7 \times 7$ | $15 \times 15$ |
| $H$ | Hidden channels | 5 | 5 |
| - | Hidden neurons | 40 | 40 |
| - | Max generations to train for | 100k | 100k |
| $\frac{S}{C}$ | Samples per class | 40 | 20 |

**Table 11:** The hyperparameters for 4 class CIFAR-10. More tuning was performed to improve the performance, and the extra layer and ReLU activation only marginally improved the performance over a simpler model. The ANCA might do better with more samples per generation, because it could then learn outlier classes instead of only the common appearances of the classes. However, that proved difficult with the time-consuming optimization.

| Hyperparameter | Description | Setup 1 |
| --- | --- | --- |
| $N_{neo} \times M_{neo}$ | Size of computational substrate | $15 \times 15$ |
| $H$ | Hidden channels | 2 |
| - | Hidden neurons | (20,20) |
| - | Activation function | ReLU |
| - | Max generations to train for | 60k |
| $\frac{S}{C}$ | Samples per class | 50 |
| $\lambda$ | Output regularization scaling | 0.0 |
| $\lambda_W$ | Weight regularization scaling | 0.0001 |

## Footnotes

1 Code: https://github.com/bioAI-Oslo/column

